# Interactions of the *Trypanosoma brucei brucei* zinc-finger-domain protein ZC3H28

**DOI:** 10.1101/2021.08.09.455650

**Authors:** Tania Bishola, Christine Clayton

## Abstract

In *Trypanosoma brucei* and related Kinetoplastids, regulation of gene expression occurs mostly post-transcriptionally, and RNA-binding proteins play a critical role in the regulation of mRNA and protein abundance. *Trypanosoma brucei* ZC3H28 is a 114 KDa cytoplasmic mRNA-binding protein with a single C(x)_7_C(x)_5_C(x)_s_H zinc finger at the C-terminus and numerous proline-, histine- or glutamine-rich regions. We here show that N-terminally tagged ZC3H28 copurifies ribosomes, various RNA-binding proteins, and the translation initiation complex EIF4E4/EIF4G3. ZC3H28 is preferentially associated with long RNAs that have low complexity sequences in their 3’-untranslated regions. When tethered to a reporter mRNA, ZC3H28 increased the mRNA level without a corresponding increase in protein expression; this suggests that it stabilized the reporter but at the same time suppressed its translation. Indeed, there was a clear negative correlation between ZC3H28 mRNA binding and ribosome density. After ZC3H28 depletion, the relative levels of ribosomal protein mRNAs increased while levels of some long mRNAs decreased, but there is no overall correlation between binding and RNAi effects on mRNA abundance. We speculate that ZC3H28 might be implicated in stabilizing poorly-translated mRNAs.

## Introduction

*Trypanosoma brucei* and related kinetoplastid parasites rely heavily on post-transcriptional mechanisms for control of gene expression. Control is required not just to determine steady-state expression, but also to respond to external stresses and to adapt to different environments in infected mammals and in the definitive host, the Tsetse fly. Most transcription is polycistronic: mRNA levels are determined by gene copy numbers, the efficiency of mRNA processing (via *trans* splicing and polyadenylation) and the rate of mRNA decay, and rates of protein synthesis also vary considerably. RNA-binding proteins are critical at all stages (Clayton, 2019).

In this paper we studied the role and interactions of ZC3H28 (Tb927.9.9450), a 114 KDa (1030-residue) protein with a single C(x)_7_C(x)_5_C(x)_s_H (CCCH) zinc finger at the C-terminus. The central part of the protein has multiple proline-, histidine- or glutamine-rich regions, including (H)_11_, (H)_17_, (Q)_11_ and (H)_14_ with a single interruption (Figure 1A). Unsurprisingly, Phyre2 predicts that about 70% of the protein lacks ordered secondary structure. ZC3H28 expression is not developmentally regulated. Results from a high-throughput RNAi screen indicated that ZC3H28 is essential in bloodstream forms, and during differentiation to, and early survival as, the procylic form (Alsford *et al*., 2011). High-throughput tagging localised it to the cytoplasm (Dean *et al*., 2016). It can also be cross-linked directly to mRNA (Lueong *et al*., 2016). In the “tethering” assay, we express in trypanosomes the protein of interest fused to the lambdaN peptide together with a reporter mRNA with “boxB” sequences that are bound with high affinity by the lambdaN peptide. In a high-throughput screen, ZC3H28 activated reporter expression (Erben *et al*., 2014), suggesting that it either increases translation efficiency, or stabilises the mRNA, or both. A study of proteins associated with the two poly(A) binding proteins showed that ZC3H28 was preferentially associated with PABP2 (Zoltner *et al*., 2018). ZC3H28 also copurified with the mRNA encoding variant surface glycoprotein (VSG), but not the mRNA encoding alpha tubulin (Melo do Nascimento *et al*., 2021)

**Figure 1.**
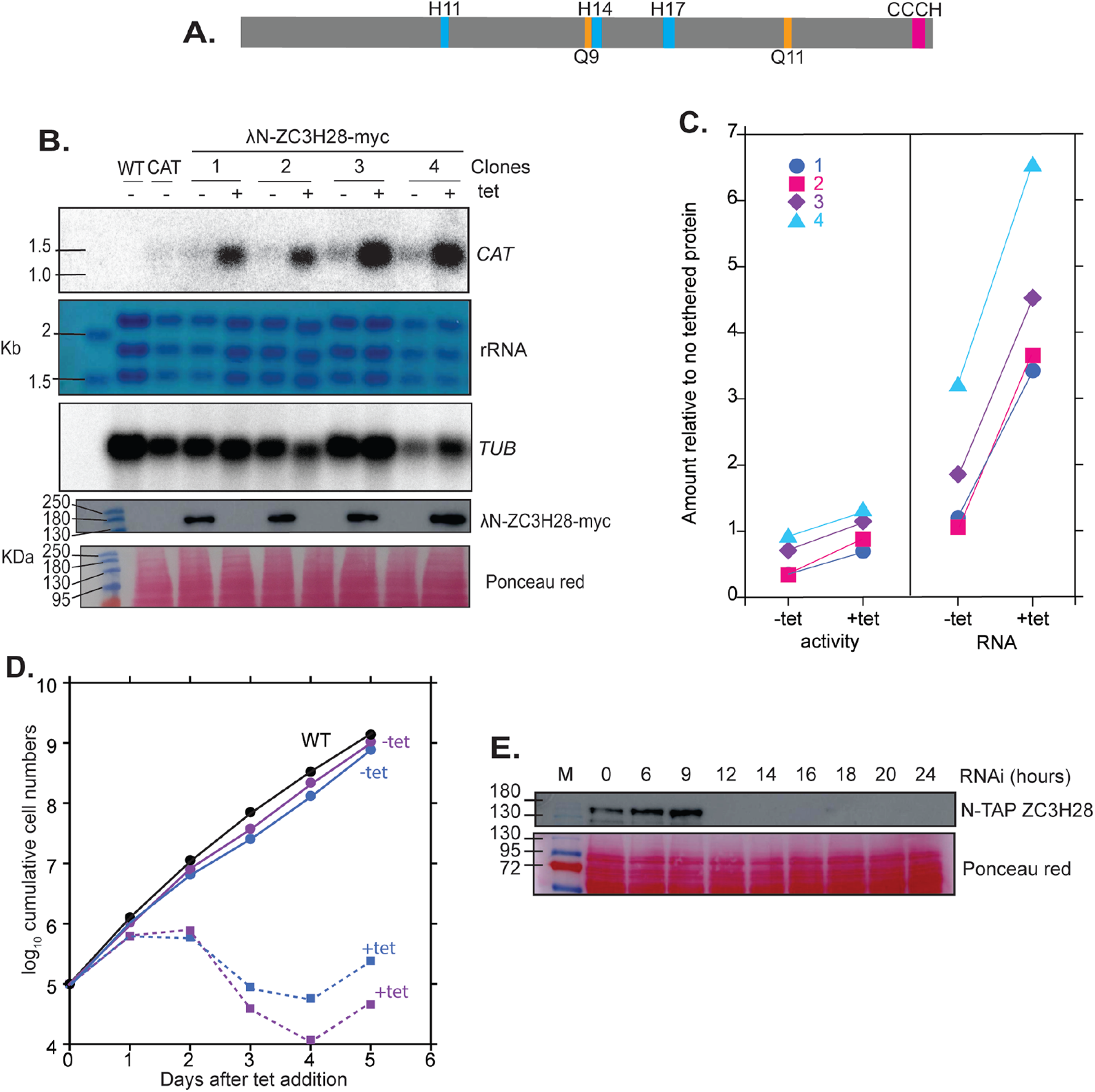
ZC3H28 is essential and increases the abundance of an attached mRNA A. Structure of ZC3H28, drawn to scale. The positions of the zinc finger domain, poly(His) and poly(Gln) sequences are indicated. B. Effects of tethering lambdaN-tagged ZC3H28 on a boxB-bearing CAT reporter mRNA. This panel shows expression of the *CAT* mRNA, with a beta-tubulin control, in the upper three panels; the lower two panels show expression of the lambdaN-tagged protein, which has a myc tag at the C-terminus. C. Effects of tethering on CAT activity (left) and mRNA (right). Results are expressed relative to cells without any tethered protein construct. D. Effect of ZC3H28 depletion on cell numbers; results for two clones are shown. E. Depletion of TAP-tagged ZC3H28 after RNAi. The stained membrane serves as the control.

One known mechanism for post-transcriptional activation in trypanosomes is recruitment of a complex containing MKT1, PBP1 (poly(A)-binding-protein-binding-protein), XAC1 (expression activator 1), LSM12 and the poly(A) binding protein PABP2 (Nascimento *et al*., 2020; Singh *et al*., 2014). The activation is likely to be a consequence of PABP2 binding and perhaps also recruitment of the translation initiation complex EIF4E6/EIF4G5. ZC3H28 was strongly and significantly enriched after affinity purification of XAC1 (Nascimento *et al*., 2020), but was barely detectable in a purified MKT1 preparation (Singh *et al*., 2014). ZC3H28 lacks a known H(N/D)PY consensus motif (Singh *et al*., 2014) for MKT1 interaction. However several proteins that enrich with MKT1 have polyglutamine tracts instead (Singh *et al*., 2014).

In this work we investigated the interactions of ZC3H28 with mRNAs and proteins, and the effects of ZC3H28 depletion on the transcriptome.

## Results

### ZC3H28 is present in kinetoplastids and a bodonid

Examination of representative Kinetoplastid genomes revealed ZC3H28 in all of them. The position in the genome upstream of the gene encoding peroxisomal protein PEX13 is also conserved although the synteny is not annotated. We also found a homologue in the bodonid *Bodo saltan*s, but not in *Euglena gracilis*. All of the sequences examined (except the *Leptomonas seymouri* sequence, which has a frame-shift near the C-terminus) have the C-terminal zinc finger and various histidine-and glutamine-rich regions (Supplementary Figure S1).

### ZC3H28 is an essential protein

Results obtained in high-throughput screens are not always reliable, so we first confirmed that lambdaN-ZC3H28 indeed activates expression of a boxB-containing chloramphenicol acetyltransferase (CAT)-encoding mRNA. Results for four independent clones confirmed roughly 2-fold increases in CAT protein but 4-6-fold increases in the amount of mRNA (Figure 1B, C). These results suggest that ZC3H28 might increase mRNA stability while suppressing translation.

Next we assessed the effects of ZC3H28 depletion on the proliferation of bloodstream-form trypanosomes. We first integrated a plasmid for inducible RNAi in EATRO1125 cells, which are competent for differentiation into the procyclic (tsetse midgut) form. *ZC3H28* RNAi only somewhat inhibited cell proliferation. We therefore instead used a cell line that we had generated for affinity purification of ZC3H28. This line was made with Lister 427 bloodstream forms, which grow to higher densities but are unable to complete differentiation. We integrated a sequence encoding a tandem affinity purification (TAP) tag in-frame with one allele, which should result in production of N-terminally tagged ZC3H28 (TAP-ZC3H28). RNAi in this line resulted in loss of the protein within 12h, and strong growth inhibition, confirming that ZC3H28 is essential in bloodstream forms (Figure 1D, E). After a few days the cells resumed growth, presumably because of protein re-expression.

It is important to note that we were unable to delete the unmodified ZC3H28 gene in the tagged cell line. This may mean that TAP-ZC3H28 is not fully functional, which in turn might have exacerbated the effect of the RNAi. Alternatively, it may be that the cells are unable to grow for long periods with only a single copy of the gene, whether the protein is tagged or not.

### Interactions of ZC3H28 with other proteins

From published resutls, it isn’t clear whether ZC3H28 is specifically associated with the MKT1 complex. We therefore examined interactions of ZC3H28 with the different components using the yeast 2-hybrid assay (Figure 2A). In this assay, ZC3H28 interacted with itself, and with PBP1, but not with MKT1. The N-terminal 527 residues (containing two poly(His) and one poly(Gln) segments) also interacted with itself and with PBP1; somewhat oddly, the N-terminal part of ZC3H28 interacted with full-length ZC3H28 only when the latter was in the “prey” configuration. Overall these results result suggest that ZC3H28 can interact-probably directly-with PBP1.

**Figure 2.**
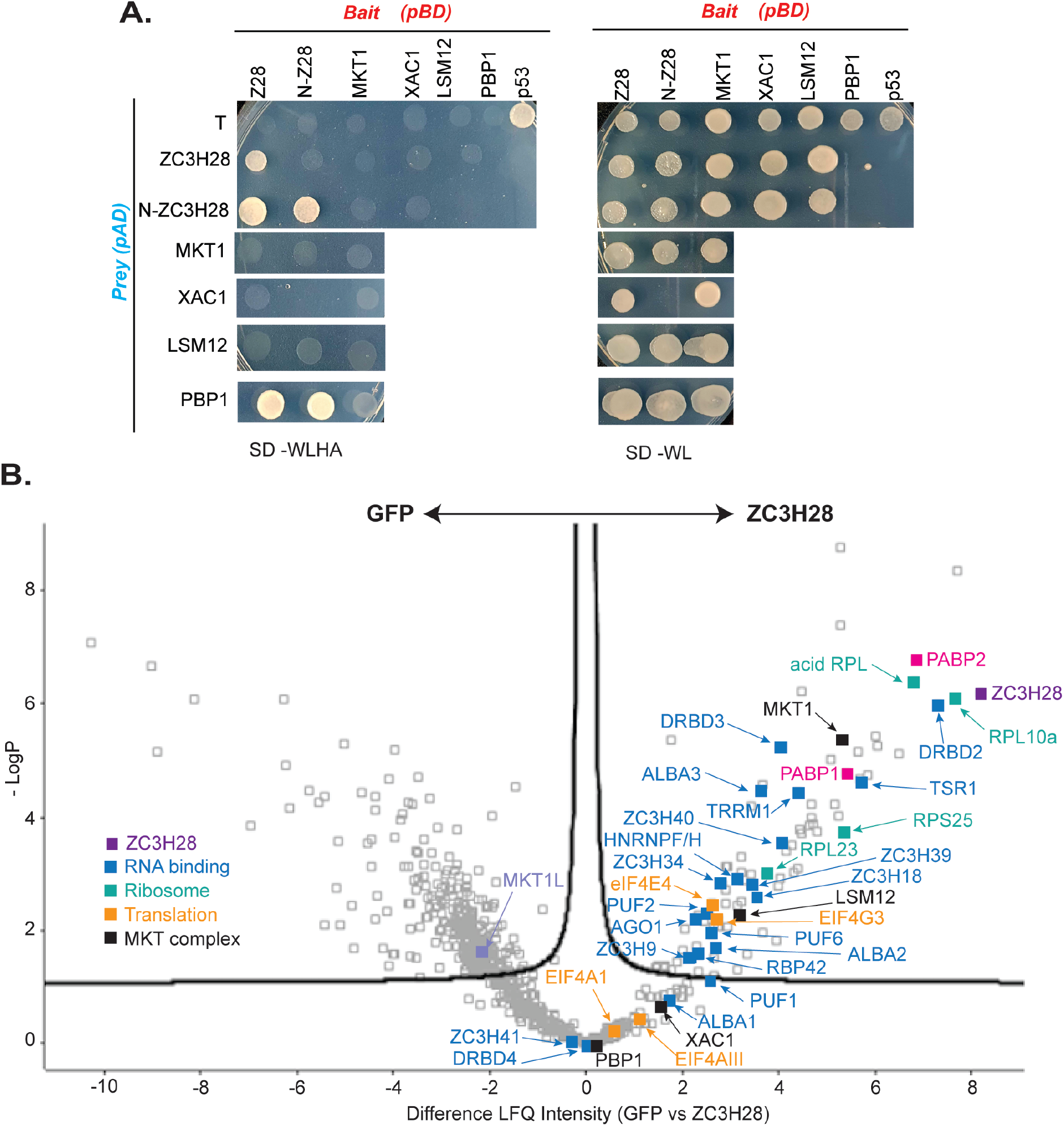
Protein interactions of ZC3H28 A. Interactions of ZC3H28 in the yeast 2-hybrid assay. *S. cerevisiae* AH109 was transformed with combinations of plasmids used as bait (pBD, DNA binding domain) and prey (pAD, transcription activating domain). The cells were grown on media lacking adenine, histidine, leucine and tryptophan (SD-WLAH) to test protein interactions and then in media containing adenine and histidine (SD-WL) to check expression of the bait and prey. The interaction between SV40 large T antigen (T) and p53 serves as positive control, and encoding lamin C (LamC) and T serves as negative control. PBP1 and XAC fused with the DNA-binding domain both activate when expressed alone (Nascimento *et al*., 2020), therefore they are used here only as activation domain fusions, which give appropriate negative control results (Nascimento *et al*., 2020). B. Volcano plot for mass spectrometry of triplicate purifications of TAP-ZC3H28, compared with TAP-GFP. The Figure was generated using Perseus software (Tyanova *et al*., 2016).

To find out whether ZC3H28 is associated with the MKT complex *in vivo*, we purified TAP-ZC3H28 and analyzed all co-purifying proteins by mass spectrometry (Supplementary Table S1, Figure 2B). Although MKT1, XAC1 and LSM12 consistently copurified, PBP1 was present in only 2 of the 3 replicates. Since TAP-ZC3H28 may not be fully functional, it is possible that the N-terminal tag interfered with some interactions. However the yeast two-hybrid constructs are also N-terminal fusions, so tags alone cannot explain the contradictory results. Thus the issue of association of ZC3H28 with the MKT1-PBP1 complex remains unresolved.

The mass spectrometry results were consistent with an association of ZC3H28 with ribonucleoprotein particles or polysomes including numerous RNA-binding proteins and the two poly(A) binding proteins. A large number of ribosomal proteins was present. Interestingly, ZC3H28 was specifically associated with just one of the five known EIF4E-EIF4G translation initiation complexes, EIF4E4-EIF4G3. Various splicing factors also co-purified, including the putative regulators TSR1 (Gupta *et al*., 2014) HNRNPF/H (Gupta *et al*., 2013), which are probably also associated with cytosolic mRNAs; and TRRM1 which has been implicated in transcription elongation (Banuelos *et al*., 2019; Levy *et al*., 2015; Naguleswaran *et al*., 2015). Considering just proteins present in all three ZC3H28 purifications, the RNA-binding proteins ALBA1, ALBA2, DRBD2, ZC3H34 and ZC3H41 were also associated with both PABP1 and PABP2; HNRNPF/H,TSR1, TRRM1, ZC3H39 and ZC3H40 were associated with PABP2, and ALBA3 with PABP1 (Zoltner *et al*., 2018). The protein of unknown function encoded by Tb927.10.9330, associated with ZC3H28, was also found with both PABPs (Zoltner *et al*., 2018). ZC3H28 also pulled down AGO1 (Shi *et al*., 2004a; Shi *et al*., 2004b), which does not copurify with the PABPs. RNA-binding proteins that were specific to the ZC3H28 purification were PUF2 (Jha *et al*., 2014), RBP42 (Das *et al*., 2012), ZC3H18 (Benz *et al*., 2011) and ZC3H9.

Overall these results would be consistent with ZC3H28 being implicated in a range of mRNA-related processes.

### mRNA interactions of ZC3H28

To find out which mRNAs are preferentially bound by ZC3H28, we purified the TAP-tagged protein, cleaved the tag with tobacco etch virus (TEV) protease and identified the co-purifying RNAs (Supplementary Table S2). The mRNA encoding ZC3H28 was 2.3-4.5-fold enriched, consistent with some purification via the nascent polypeptide. Unexpectedly, *VSG* mRNA was not preferentially bound (Supplementary Table S2). In contrast, ZC3H28 preferentially associated with long mRNAs, especially those with long 3’-untranslated regions (Figure 3A), although the correlation was only partial, suggesting a degree of sequence specificity (Figure 3B). Consistent with the length bias, the list of 180 mRNAs that were at least 3-fold enriched in the bound fraction included 13 mRNAs encoding protein kinases and 12 encoding RNA-binding proteins, which tend to be long mRNAs; and there was no association with mRNAs encoding ribosomal proteins, which are mostly very short (Clayton, 2019; Erben *et al*., 2021) (Figure 3C). A search for motifs using >3x-enriched transcripts, with a set of length-matched controls with average binding ratios of less than 1, revealed weak enrichment of poly(U) and poly(AU) sequences. These are common in trypanosome 3’-UTRs and it turned out that the median 3’-UTR lengths of the length-matched controls were half those for the 3xbound set. We therefore looked for motifs in the 3’-UTRs only, again using appropriately length-matched controls. This revealed significant enrichment of poly(AU) and polypurine tracts in the bound mRNA 3’-UTRs (Figure 3C). However, none of the motifs was present in all bound mRNAs, or exclusive to them. For example, the sequence “AUAUAUAUA” is present at least once in 49 of the 106 bound 3’-UTRs but also in 36 of the 103 unbound ones. The basis for the selectivity of ZC3H28 is therefore not clear.

**Figure 3.**
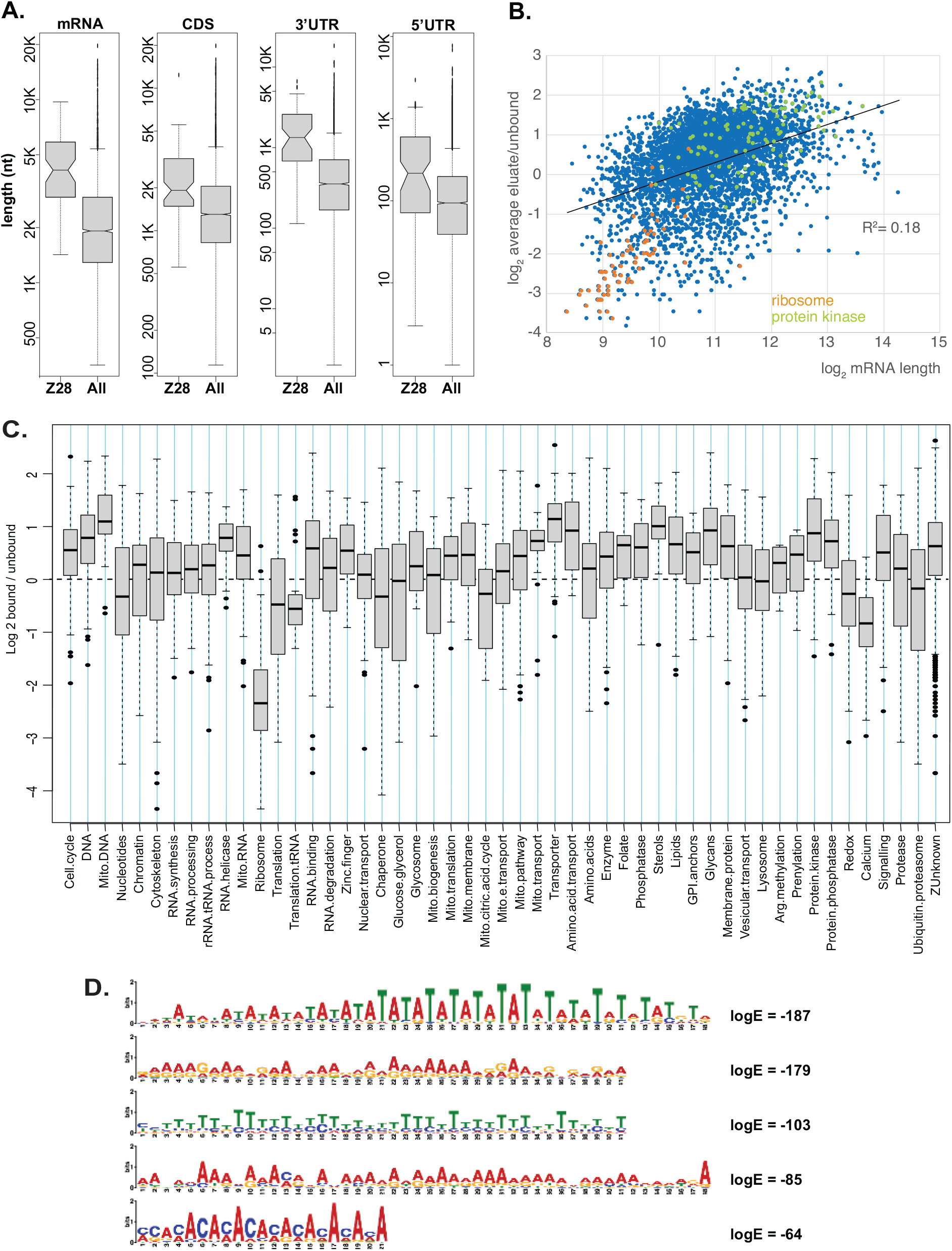
Interactions of ZC3H28 with mRNA. A. Characteristics of mRNAs that were at least 3-fold enriched with TAP-ZC3H28 relative to the unbound fraction. Results for the whole transcriptome are shown for comparison. All results in this Figure are for the set of unique genes, to avoid over-counting repeated genes. B. Plot for the whole dataset: enrichment with ZC3H28 relative to mRNA length. C. The set of unique genes was placed in functional categories (Supplementary Table S4) and binding to ZC3H28 was plotted for each category. The dotted line indicates equal distribution in both fractions. D. Motifs enriched in the 3’-UTRs of ZC3H28-bound mRNAs relative to size-matched controls showing less than 1-fold average enrichment in the bound fraction.

### The effect of ZC3H28 depletion on the transcriptome

The results so far would all be consistent with partially selective association of ZC3H28 with translating mRNAs. From the tethering result we would expect association of ZC3H28 to enhance expression, particularly at the mRNA level. To test the role of ZC3H28 further we examined the effect of ZC3H28 depletion on the transcriptome (Supplementary Tables S3 and S4). First, we looked at triplicate samples from cells grown without tetracycline, and after 10h incubation with tetracycline. This time point was chosen because 10h RNAi induction was insufficient to affect growth (Supplementary Figure S2). Very few differences were found, so we also examined further duplicate samples after 14h or 16h with tetracycline, and 24h either with or without tetracycline. Throughout this period the cells with tetracycline grew only very slightly slower than the controls, but after 24h the numbers started to decrease (supplementary Figure S2). Intriguingly, a principal component analysis (Figure 4A) clearly separated the first set of controls (labelled -tet 1, -tet 2, and -tet3) from the second set (labelled A-tet and B-tet). The 10h induction (+tet 10h) clustered with its own control (Figure 4A), while the 24h induction (A-24h, B-24h) was quite similar to its control. In contrast, RNAi induction for 14h and 16h had very clear effects on the transcriptome.

**Figure 4.**
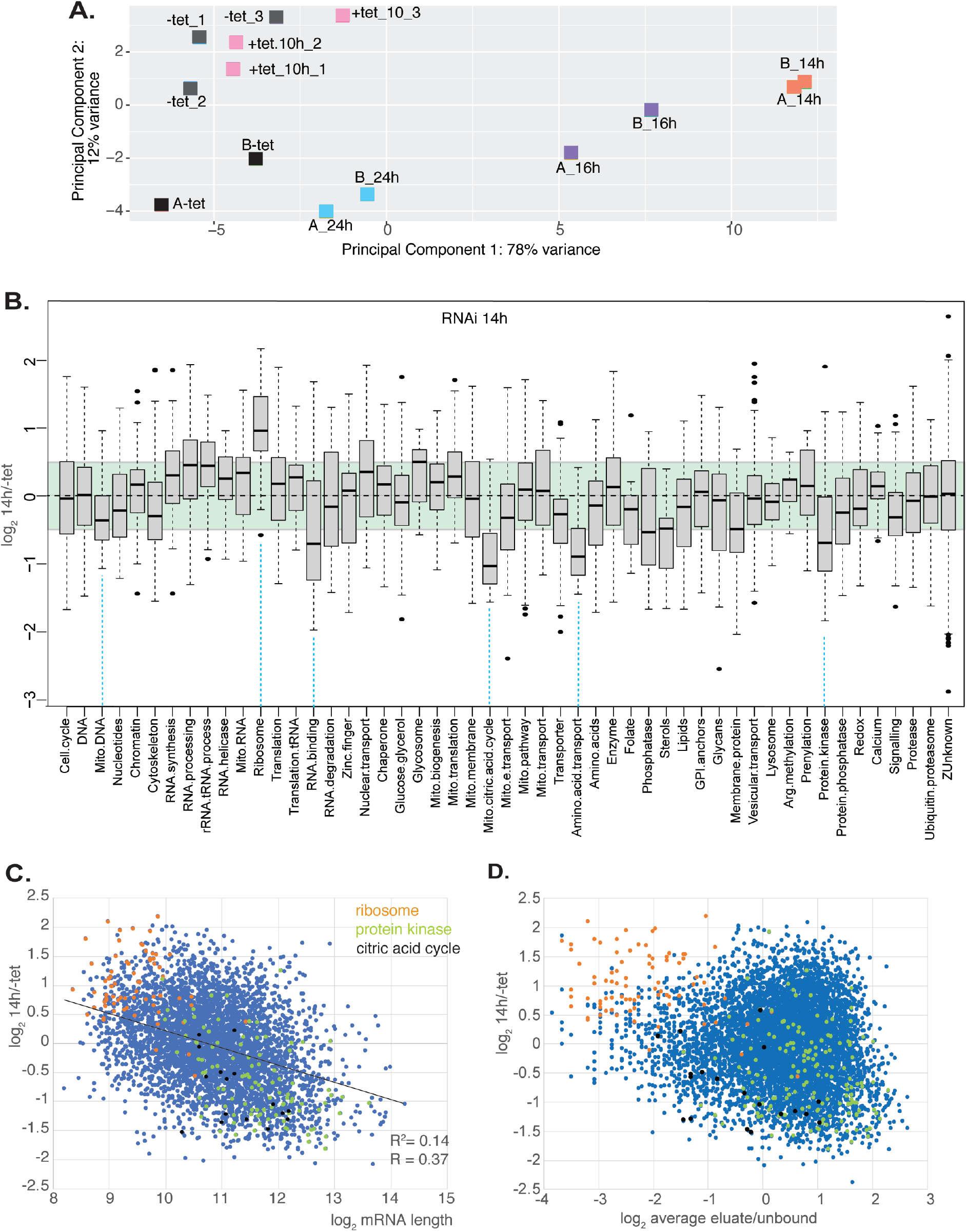
Effects of ZC3H28 depletion on the transcriptome A. Principal component analysis. Samples without induction are in black and grey; others are colour-coded according to the time with tetracycline. All results in this Figure are for the set of unique genes, to avoid over-counting repeated genes. B. The set of unique genes was placed in functional categories (Supplementary Table S4) and the effects of ZC3H28 depletion (14h RNAi) were plotted for each category. The green background is the median and 25th percentiles for the total dataset. C. The effect of RNAi (y-axis) was plotted against the annotated mRNA length (x-axis). Only mRNAs with two annotated untranslated regions were included. Correlation coefficients were calculated in Microsoft Excel. D. Lack of relationship between the ZC3H28 RNAi effect (y-axis) and ZC3H28 binding (x-axis).

These results were strange so we first looked at the difference between the two sets of controls. The cell densities for the triplicate samples were 5-6 × 10^5^/ml, whereas the duplicates were harvested at approximately 1×10^6^/ml. The mRNAs that were 1.5-fold significantly (Padj <0.05) increased at the higher density included those encoding the procyclins and some proteins of mitochondrial metabolism, suggesting changes consistent with very early differentiation (Supplementary Table S4). Decreases of mRNAs encoding, for example, RNA polymerase I subunits, nucleotide transporters and translation elongation factor 2 would be consistent with slowing growth-although mRNAs encoding various RNA processing proteins, polymerase II subunits and flagellar proteins were increased. After 24h of RNAi induction the cell densities were 1.3-1.6 × 10^6^/ml; perhaps this suppressed the RNAi effect. Amounts of total mRNA (Supplementary Figure S3A) and overall protein synthesis were not decreased by *ZC3H28* RNAi (Supplementary Figure S3B), but the rate of protein synthesis was reproducibly lower at densities above 1 x10^6^/ml (Supplementary Figure S3B) both with or without RNAi.

The effects of ZC3H28 RNAi on the transcriptome peaked at about 14 hours after tetracycline addition (Figure 4A). More than 500 mRNAs were two-fold significantly increased, and nearly 700 decreased, at 14h; after 16h this had decreased to 101 and 243, respectively. We therefore concentrated on the effects after 14h. The strongest effects were clearly on mRNAs encoding ribosomal proteins (Figure 4B): 48% were more than 2-fold increased and 77% more than 1.5-fold. In contrast mRNAs encoding RNA-binding proteins, protein kinases, citric acid cycle enzymes and amino acid transporters were relatively decreased (Figure 4B). The specificity here was unclear, since the effects were weakly inversely correlated with mRNA length (Figure 4C). (The precise correlation is unknown because many 3’-UTRs are annotated to be shorter than sequence coverage suggests.) Shorter RNAs, which tended not to be bound by ZC3H28, also tended to increase after RNAi, and longer RNAs, which tended to be bound, decreased. The 39 mRNAs that were both at least 3-fold enriched with TAP-ZC3H28, and 2-fold decreased after RNAi, had a median annotated length of 4.8 kb; their products included 7 protein kinases and five RNA-binding proteins. In general, effects at 16h were similar but slightly less pronounced.

Overall, there was no correlation between the degree of RNA binding and effects after RNAi (Figure 4D). However the mRNAs that were at least 3-fold enriched did have a 1.6-fold median decrease after loss of ZC3H28, which is consistent with the ability of ZC3H28 to increase mRNA levels in the tethering assay.

We finally looked for additional correlations. The bound mRNAs are mostly not cell-cycle regulated (Archer *et al*., 2011) and there was no correlation with developmental regulation of either mRNA level (Fadda *et al*., 2014) or translation (Antwi *et al*., 2016). Interestingly, however, binding of mRNAs to ZC3H28 was clearly negatively correlated with the ribosome density on the coding region (Antwi *et al*., 2016) (Figure 5A). This appears not to be secondary to other mRNA characteristics, since there is no correlation between ribosome density on the coding region and mRNA length or 3’-UTR length (Supplementary Table S4). There was also no correlation between the effects of RNAi and either ribosome density or mRNA half-life, and no correlation either between ZC3H28 mRNA binding and mRNA half-life. This result, combined with that from tethering, suggests that ZC3H28 might act to protect poorly-translated mRNAs from degradation.

**Figure 5.**
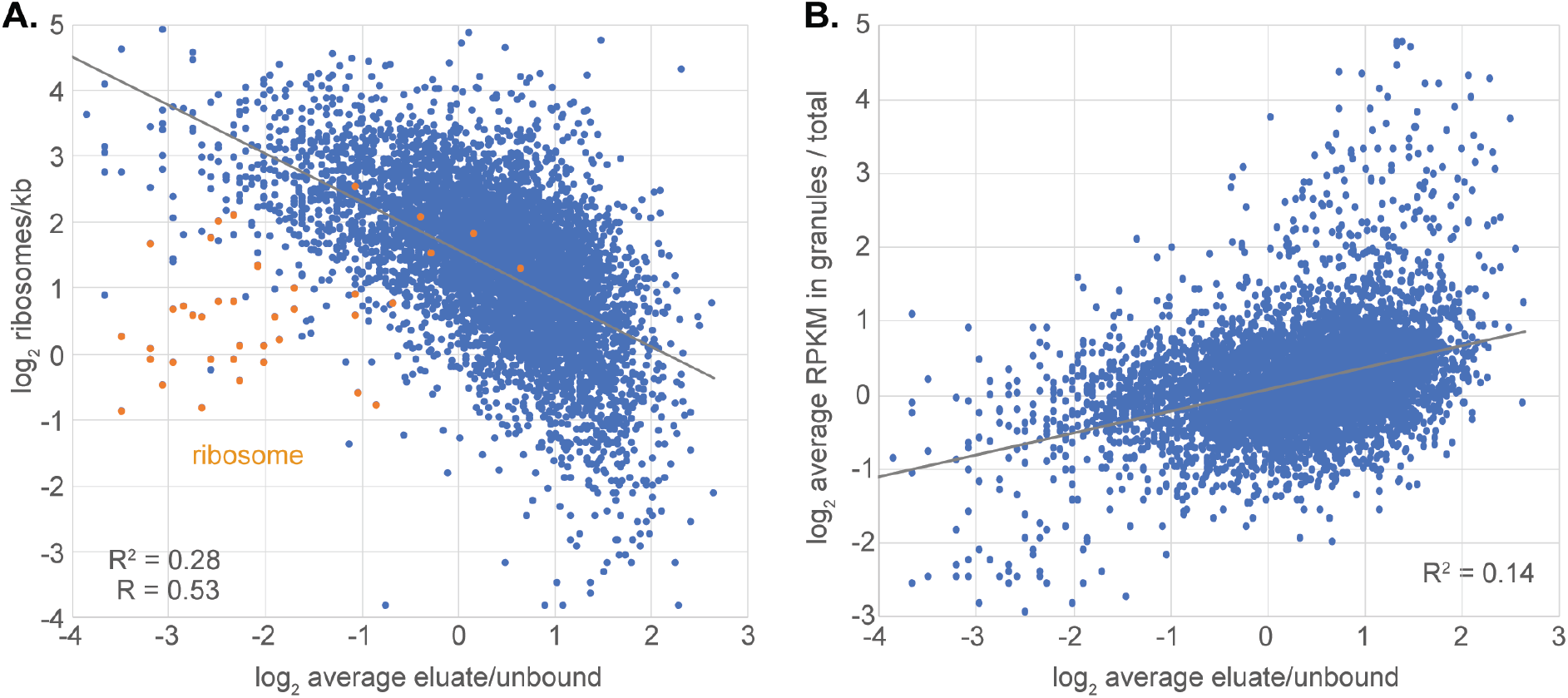
ZC3H28 and translation A. Binding of mRNAs to ZC3H28 is on the x-axis, and the ribosome density on the y-axis. B. Binding of mRNAs to ZC3H28 is on the x-axis, and enrichment in procyclic-form starvation granules (Fritz *et al*., 2015) is on the y-axis.

## Discussion

Our cumulative results suggest that ZC3H28 acts to decrease translation of preferentially associated mRNAs. ZC3H28 has only a single zinc finger. A crystal structure of the two CCCH domains of TIS11 showed that each CCCH domain could bind specifically to four nucleotides (Hudson *et al*., 2004), and that the region 5’ to the zinc finger contributed to specificity. With only a single zinc finger, not much specificity would be expected unless ZC3H28 acts as a multimer. We found that bound mRNAs tended to be relatively long, especially in the 3’-untranslated region, with long low-complexity sequences. Several different low complexity sequences were seen, and there was no discernable base specificity.

The effects of ZC3H28 depletion on the transcriptome where quite surprising. First, we found clear indications that non-depleted cells at a density of about 1 x10^6^/ml were already starting to undergo changes typical of differentiation to the stumpy form, although these were Lister 427 parasites which are incapable of growth arrest and stumpy differentiation. These cells also had a somewhat lower translation rate than cells at 5 x10^5^/ml. After 14h of RNAi induction - which corresponds to less than 5h without ZC3H28 (Figure 1 E), depleted cells (at densities around 7×10^5^/ml) showed significant effects on over 1000 mRNAs, especially preferential loss of longer transcripts. Many of these effects could be indirect since most of the decreased mRNAs were not actually bound by tagged ZC3H28. However only 2h later the differences had begun to diminish and after 24h, at 1 x10^6^/ml, the transcriptomes had reverted to be similar to those of controls. These results suggest that ZC3H28 is most important in growing parasites.

As is the case for many experiments with mRNA-binding proteins, none of the correlations is very strong, because the final status of mRNAs will be determined not by ZC3H28 alone, but by the additional actions of all the other regulatory molecules.

Tethering of ZC3H28 to a reporter increased the amount of mRNA without a corresponding increase in protein, suggesting that translation was suppressed. Consistent with this, ZC3H28 is preferentially associated with mRNAs that have low ribosome occupancy in their coding regions. A possible mechanism for the two effects could be sequestration of mRNAs in non-translating compartments. Intriguingly, C-terminally GFP-tagged ZC3H28 was found in cytoplasmic aggregates in slightly starvation-stressed procyclic forms (see http://tryptag.org/?query=Tb927.9.9450), although it was not enriched in granules purified after more prolonged starvation (Fritz *et al*., 2015). Aggregation is expected to occur in proteins with long poly(His) or poly(Gln) tracts, as seen in ZC3H28, and unsurprisingly, in the yeast two-hybrid system, ZC3H28 showed self-interaction. Intriguingly, mRNAs that are found in procyclic granules did show preferential association with ZC3H28 in bloodstream forms (Figure 5B). Overall, our results suggest that ZC3H28 is implicated in controlling translation of a subset of mRNAs in growing trypanosomes.

## Materials and Methods

### Trypanosome culture and modification

The experiments in this study were carried out using monomorphic *T. brucei* Lister 427 bloodstream form parasites constitutively expressing the Tet-repressor (Alibu et al. 2005). The parasites were cultured at 37°C as routinely in HMI-9 medium supplemented with 10% heat inactivated fetal bovine serum (v/v), 1% (v/v) penicillin/streptomycin solution (Labochem international, Germany), 15 µM L-cysteine, and 0.2 mM β-mercaptoethanol in the presence of 5% CO2 and 95% humidity (Hirumi and Hirumi, 1989). During proliferation, the cells were diluted to 1×10^5^ cells/ml and maintained in density between 0.2-2×10^6^ as described in (Clayton, 1999). Cell densities were determined using a Neubauer chamber. For generation of stable cell lines, ∼1-2 × 10^7^ cells were transfected by electroporation with 10 µg of linearized plasmid at 1.5 kV on an AMAXA Nucleofector. Selection of newly transfectants was done after addition of appropriate antibiotic at the following concentrations: 1 μg/ml puromycin, 2.5 μg/ml phleomycin (InvivoGen), 5 μg/ml hygromycin B (Calbio-chem. 10 μg/ml blasticidin (InvivoGen). Independent clones were obtained by serial dilution.

### Genetic manipulation of trypanosomes

A cell line with *in-situ* TAP-ZC3H28 gene was generated by replacing one endogenous copy of ZC3H28 with a gene encoding N-terminally TAP tagged ZC3H28. For this purpose, a construct with puromycin resistance gene plus a TAP tag cassette was flanked on the 5’-end with a fragment of ZC3H28 5’-UTR. Also, downstream on the 3’-end, the N terminal region of ZC3H28 ORF was cloned in frame with the TAP tag. Prior to transfection, the plasmid (pHD3236) was cut with SacI and ApaI enzymes to allow homologous recombination. Using the cell lines expressing the *in-situ* N-TAP ZC3H28, we were unable to knock-out the other copy of ZC3H28. Gene fragments for RNAi were selected based on default settings of the RNAit software (Redmond *et al*., 2003) and cloned into the pHD 1146 plasmid. For the tethering assays, cell lines constitutively expressing the CAT reporter with boxB and the actin 3’-UTR were co-transfected with plasmids encoding the ZC3H28 in fusion with the λN-peptide and a myc tag (Erben *et al*., 2014). The primers and plasmids used are listed in Supplementary Table S5.

### DNA extraction

Genomic DNA from *T. brucei* was isolated using 1-2 × 10^8^ cells as follows. The cell pellet was collected by centrifugation (2300 rpm, 8 minutes), washed once in cold 1x PBS, and lysed in 0.5 ml of EB buffer (10 mM Tris-HCl pH 8.0, 10 mM NaCl, 10 mM EDTA). RNA was digested with addition of 12 µl RNAse A (1mg/ml stock solution, Sigma-Aldrich) at 37°C for 30 minutes. Proteins were precipitated using 200 µl ice-cold 5M ammonium acetate followed by centrifugation at maximum speed for 5 minutes. The supernatant containing the DNA was transferred to a new tube. The DNA was then precipitated with 0.7x isopropanol followed by centrifugation at maximum speed for 15 minutes. The pellet was then washed once with 75 % ethanol to remove salts and then again with 100 % ethanol followed by centrifugation for 5 minutes. The DNA pellet was then dried for approximately 5 minutes and dissolved in TE buffer (10 mM Tris pH 7.5, 1 mM EDTA pH 8.0) at 37°C. The concentration was measured using a Nanodrop. PCR was done using Taq Polymerase according to the manufacturer’s instructions (New England Biolabs).

### RNA manipulation

To identify RNAs bound to ZC3H28, approximately 1×10^9^ cells expressing *in-situ* N-TAP tagged ZC3H28 with a concentration of 1×10^6^ cells/ml were pelleted by centrifugation at 3000 rpm for 13 minutes at 4°C. The pellet was washed twice in cold 1X PBS and collected by centrifugation at 2300 rpm for 8 minutes at 4°C and then snap frozen in liquid nitrogen. The RNA immunoprecipitation was done essentially as described in (Mugo & Clayton, 2017). The cell pellet was lysed in 1 ml of the lysis buffer (20 mM Tris pH 7.5, 5 mM MgCL_2_, 0.1% IGEPAL, 1 mM DTT, 100 U RNAsin, 10 μg/ml leupeptin, 10 μg/ml Aprotinin) by passing 20 times through a 21G x ½ needle using a 1 ml syringe and 20 times through a 27G x ¾ needle using a 1 ml syringe. The lysate was cleared by centrifugation at 15,000 g for 15 minutes at 4°C and the supernatant was transferred to a new tube. The salt concentration was then adjusted to 150 mM KCl. The cell extracts were afterwards incubated with 40 μl of IgG-coupled magnetic beads (Dynabeads™ M-280 Tosylactivated, Invitrogen) for 3 hours at 4°C and the flow-through (unbound) fraction was collected by magnetic separation as the negative control. peqGOLD TriFast™ FL REAGENT was added to the unbound fractions and kept at −80°C for further RNA extraction. After three washing with IP buffer (20 mM Tris pH 7.5, 5 mM MgCL_2_, 150 mM KCl, 0.1% IGEPAL, 1 mM DTT, 100 U RNAsin, 10 μg/ml leupeptin, 10 μg/ml Aprotinin), the tagged protein was eluted from beads using 150 units of TEV protease at 4°C for overnight. The eluate was transferred to a fresh tube, two volumes of peqGOLD TriFast™ FL reagent were added, and samples were stored at −80 °C until further processing. RNA was isolated from released and bound fractions according the manufacturer’s instructions. Total RNA from the unbound and the eluate fraction were depleted of ribosomal RNA (rRNA) using RNAse H and a cocktail of 131 DNA oligos (50 bases) complementary to the trypanosome rRNAs. The rRNAs hybridized to the oligonucleotides were digested with RNAse H (NEB, M0297S) as previously described in (Minia *et al*., 2016). Following rRNA depletion, the samples were subjected to DNAse I treatment in order to remove any trace of oligonucleotides using the Turbo™ DNAse kit (Invitrogen, ThermoScientific). The RNA samples were afterward purified using the RNA Clean & Concentrator™ −5 kit (ZYMO RESEARCH) following the manufacturer’s instructions. The recovered purified RNA from both bound and unbound samples were then analyzed by RNA-Seq.

The purified RNA (5-10 µg) was mixed with 2X RNA loading dye (1,6X MOPS buffer, 7% formaldehyde, 65% formamide, 50 µg/ml ethidium bromide, 0.025% bromophenol blue), denatured for 10 minutes at 65°C and then resolved on formaldehyde agarose gel. The RNA was afterwards blotted onto Nylon membranes (Amersham Hybond-N+, GE Healthcare, RPN203B) with 10X saline-sodium citrate buffer (SSC) by capillary transfer overnight. The RNA was then cross-linked to positively charged membranes using a UV-crosslinker (Stratagene UV Stratalinker 2400, 2×240 mJoules) and stained with methylene blue (SERVA) for 10 minutes. The northern blots were pre-hybridized in hybridization solution for 1h at 65°C and then hybridized in the same solution with the appropriate (α-^32^P) dCTP radioactively labelled DNA probes from CAT and tubulin genes for overnight at 65°C. Labelling the DNA probes was done with Prime-IT RmT Random Primer Labelling Kit, Stratagene. The following day, the blot was washed twice for 10 minutes at room temperature with wash solution 1 (2x SSC, 0.1% SDS), twice for 10 minutes with wash solution 2 (1x SSC buffer, 0.1% SDS) and twice for 10 minutes at 65°C with wash solution 3 (0.1x SSC, 0.1% SDS). For spliced leader detection, a 39-mer oligonucleotide complementary to the spliced leader was labelled with [γ32P]-ATP using T4 polynucleotide kinase (NEB) and incubated with the membrane overnight at 42°C. Afterwards, the blots were exposed to autoradiography films and the signals were detected with the phosphorimager (Fuji, FLA7000). The images were processed and quantified using ImageJ.

### RNA sequencing and data analysis

RNA sequencing was done at the CellNetworks Deep Sequencing Core Facility at the University of Heidelberg. NEBNext Ultra RNA Library Prep Kit for Illumina (New England BioLabs Inc.) was used for library preparation. The libraries were multiplexed (6 samples per lane) and sequenced with a Nextseq 550 system, generating 75 bp single-end sequencing reads. This was done using a custom pipeline (Leiss *et al*., 2016) that incorporated the following steps. Before analysis, the quality of the raw sequencing data was checked using FastQC (http://www.bioinformatics.babraham.ac.uk/projects/fastqc). Cutadapt (Martin, 2011) was used to remove sequencing primers, poly(A) tails and spliced leaders. After primer removal, the sequencing data were aligned to *T. brucei* 927 reference genome using Bowtie2 (Langmead & Salzberg, 2012), allowing 1 alignment per read, then sorted and indexed using SAMtools (Li *et al*., 2009). Reads aligning to open reading frames, annotated 3’-untranslated regions, and functional non-coding RNAs were counted. The alignment and counting were then repeated with Lister 427 genome (2018 assembly) (Müller *et al*., 2018). For comparative and enrichment analyses, we used a list of unique genes modified from (Siegel *et al*., 2010) in order to avoid giving excessive weight to repeated genes and multigene families. For the RIP-Seq, the reads per millions were counted and the ratios of eluate versus unbound were calculated. An mRNA was considered as “bound mRNA” if the lowest ratio was at least 3. The motif enrichment search was done using MEME (Bailey, 2011). Annotated 3’-UTRs were downloaded from TritrypDB. Analysis of differentially expressed genes after ZC3H28 RNAi was done in R using the DESeqUI (Leiss & Clayton, 2016), a customized version of DESeq2 package (Love *et al*., 2014) adapted for trypanosome transcriptomes. Statistical analyses were done using R and Microsoft Excel.

### Yeast two-hybrid assays

The Matchmaker Yeast two-hybrid system (Clontech) was used to test direct protein-protein interactions according to the manufacturer’s instructions. The coding sequence of ZC3H28 was PCR-amplified from genomic DNA and cloned into pGBKT7 and pGADT7 plasmids. The prey and the baits plasmids of ZC3H28 as well as those of the MKT1complex proteins were co-transformed pairwise into AH109 yeast strains. Selection was done initially on double drop-out (DDO) plates (i.e., SD medium lacking Tryptophan and Leucine) to check expression of both bait and prey. The growth was then checked on quadruple drop-out (QDO) plates (i.e., lacking Tryptophan, Leucin, Histidine and Adenine) that indicates positive interactions. The interaction between p53 and SV40 large T-antigen and the combination of LaminC and SV40 large T antigen served as positive and negative controls, respectively.

### Protein purification and mass spectrometry analysis

Approximately 1×10^9^ cells expressing either *in-situ* N-TAP ZC3H28 (pHD3236) or tet-inducible GFP-TAP (pHD1743) with a concentration of 1×10^6^ cells/ml were harvested by centrifugation. For each cell line, three technical replicates were done without RNAse A treatment. The cell pellet was resuspended in 10 ml of cold 1x PBS and centrifuged at 2800 rpm for 8 minutes at 4° and the pulldowns were done as described above, except that after TEV cleavage, the eluate was collected by magnetic separation. To remove His-tagged TEV, 10 µL of equalization buffer (200 mM sodium phosphate, 600 mM sodium chloride, 0.1 % Tween-20, 60 mM imidazole, pH 8.5), as well as 30 µL of Ni-NTA-magnetic beads were added and incubated with the samples for 30 min at 20 °C while rotating. Ni-NTA magnetic beads were separated using a magnetic rack and the supernatant was collected and stored in 6X Laemmli buffer at −80 °C. Eluted proteins were separated on 12% SDS-polyacrylamide gel until the running front had migrated roughly 2 cm, thereafter the gel was stained with Coomassie blue and destained with destaining solution (10 % acetic acid, 50 % methanol in H_2_O). Two areas per lane were cut and analyzed in the ZMBH Mass Spectrometry facility via the Ultimate 3000 liquid chromatography system directly coupled to an Orbitrap Elite mass spectrometer (Thermo Fisher). MS spectra (m/z 400–1600) were acquired in the Orbitrap at 60,000 (m/z 400) resolution. Fragmentation in CID cell was performed for up to 10 precursors. MS2 spectra were acquired at rapid scan rate. Raw files were processed using MaxQuant (version 1.5.3.30; J. Cox, M. Mann, Nat Biotechnol 2008, 26, 1367) for peptide identification and quantification. MS2 spectra were searched against the TriTrypDB-8.1TREU927-AnnotatedProteins-1 database (containing 11567 sequences). Data were analyzed quantitatively and plotted using Perseus software (Version 1.6.15.0).

### Protein detection by western blotting

For western blotting, 1-5 × 10^6^ cells were collected by centrifugation at 3000 rpm for 5 minutes, washed twice in ice-cold 1x PBS, lysed in 2× Laemmli buffer and heated at 95°C for 10 minutes. The protein samples were separated using 12 % SDS-PAGE gels, and processed as described previously in Minia et al., 2016. The following antibodies were used for specific protein detection: anti-PAP (1: 1:5000, rabbit, Sigma), anti-myc (mouse, 1:1000); anti-rabbit IgG (for pull-downs). The proteins were detected by enhanced chemiluminescence according to the manufacturer’s instructions (Amersham Biosciences).

### CAT Assay

To perform the CAT assay experiment, approximately 2×10^7^ cells expressing the CAT reporter gene were harvested at 2300 rpm for 8 minutes and washed three times with 1x cold PBS. The pellet was re-suspended in 200 μl of CAT buffer (100mM Tris-HCl pH 7.8) and lysed by freeze-thawing three times using liquid nitrogen and a 37°C heating block. The supernatants were then collected by centrifugation at 15,000×g for 5 min and kept on ice. The protein concentrations were determined by Bradford assay (BioRad) according to the manufacturer’s protocol. For each setup, 0.5 μg of protein in 50 μl of CAT buffer, 10 μl of radioactive butyryl CoA (^14^C), 2 μl of chloramphenicol (stock: 40mg/ml), 200 μl of CAT buffer and 4 ml of scintillation cocktail were mixed in a Wheaton scintillation tube HDPE (neoLab #9-0149). The incorporation of radioactive acetyl group on chloramphenicol was measured using program 7 of Beckman LS 6000IC scintillation counter.

### Pulse labelling

For each time point, approximately 4 × 10^6^ cells were collected at room temperature. The pellet was washed twice with ice-cold 1x PBS followed by centrifugation at 4,000g for 3 minutes. The cell pellet was then resuspended in 400 μl labeling medium (Dulbecco’s Modified Eagle Medium (Gibco) lacking L-methionine and cysteine) at 37 °C for 1 hour. 2 μl of L-[^35^S] methionine (about 20 μCi) was then added. The cells were incubated for 1 h at 37 °C and afterwards collected by centrifugation at 4,000 g for 3 minutes at RT. The pellet was washed twice with 1x PBS and then resuspended in in 15 μl of Laemmli lysis buffer. The proteins were separated in a 12% SDS gel. The gel was the dried unto a Whatman paper, then exposed to autoradiography films and the signals were detected with the phosphorimager (Fuji, FLA7000).

The labeling medium was Dulbecco’s modified Eagle’s medium (Gibco, high-glucose, containing pyridoxine hydrochloride, lacking L-glutamine, sodium pyruvate, L-methionine, and L-cysteine), supplemented with 25 mM HEPES, 2 mM glutamine, 0.1 mM hypoxanthine, 0.0028% b-mercaptoethanol, 0.05 mM bathocupronsulfate, and 10% heat-inactivated fetal calf serum (previously dialyzed against 30 mM HEPES, pH 7.3, 150 mM NaCl).(Leiss *et al*., 2016). The transcriptome data were analysed using DeSeqU1 (Leiss & Clayton, 2016).

## Data availability

The transcriptomes are available with accession numbers E-MTAB-10674 (ZC3H28-associated RNA) and E-MTAB-10751 (effect of RNAi). The mass spectrometry proteomics data have been deposited to the ProteomeXchange Consortium via the PRIDE partner repository (Perez-Riverol *et al*., 2019) with the dataset identifier PXD027792.

## Acknowledgements

This work was partially funded by Deutsche Forschungsgemeinschaft grant number Cl112/28-1 to CC, and by core support from the state of Baden-Württemberg. We thank Claudia Helbig and Ute Leibfried for technical assistance, David Ibberson of the Bioquant sequencing facility (University of Heidelberg) for cDNA library construction and RNA sequencing. Mass spectrometry was done in the ZMBH Core Facility by Thomas Ruppert and Sabine Merker. We are indebted to Prof. Dr. Nina Papavasiliou (DKFZ, University of Heidelberg) and Prof. Dr. Luise Krauth-Siegel (BZH, University of Heidelberg) for allowing us to share their laboratories including equipment and reagents after the flood in the ZMBH. We acknowledge the support of Andrea Zanotti (AG Lemberg) for the assistance in the pulse labelling experiment.

## Author contributions

T.B. performed all the experiments. T.B. and C.C. were both involved in conceptualization, methodology, data curation, formal analysis, validation, investigation and visualization, writing-original draft, review and editing. C. C. was responsible for supervision, funding acquisition, and project administration.

## Supplements

**Supplementary Figure S1**

Alignment of protein sequences for selected kinetoplastids and *Bodo saltans*.

The alignment was done using Phyre2 with default settings. The tree at the top shows the source organisms.

**Supplementary Figure S2.**
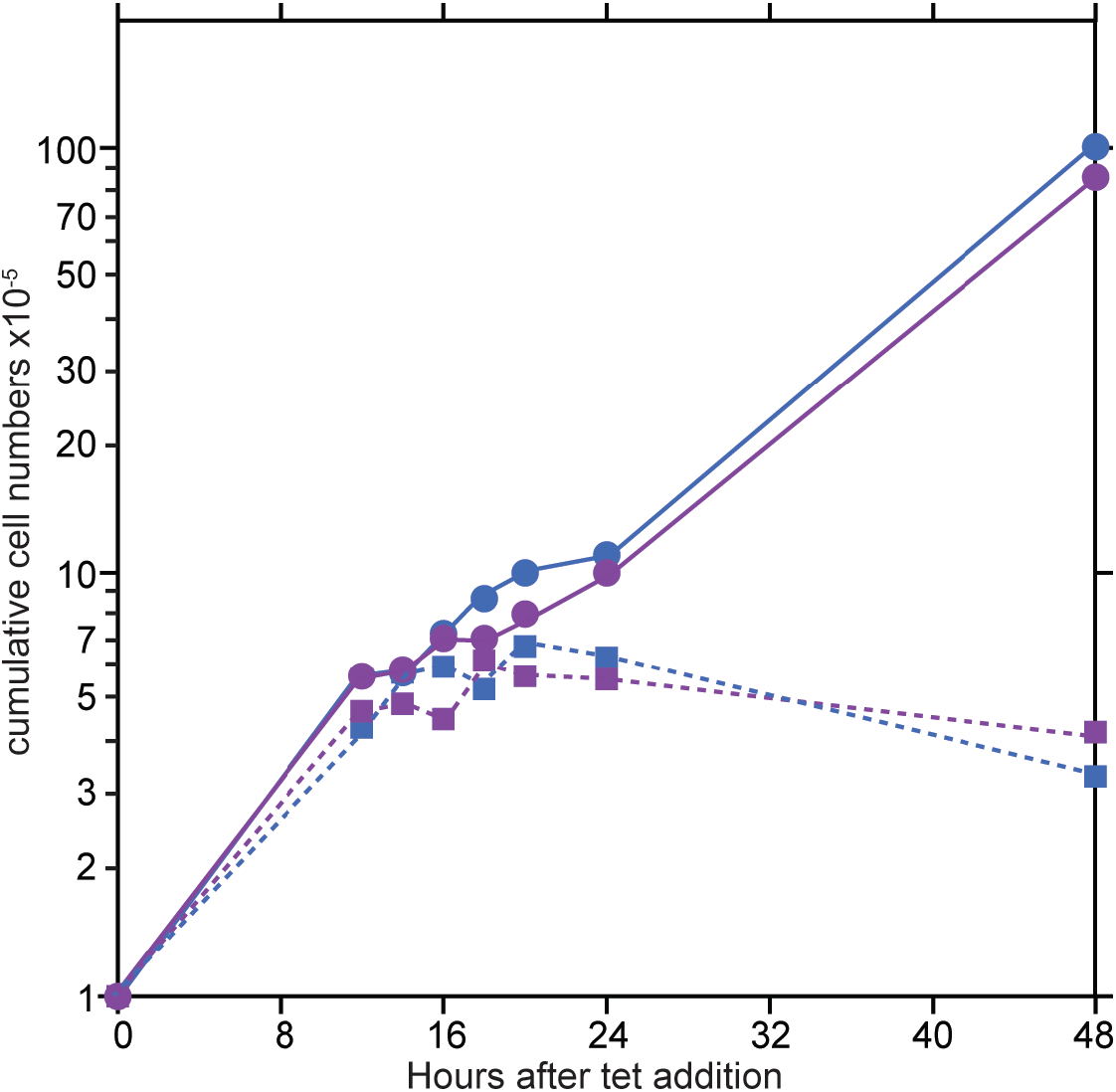
Growth of trypanosomes used for RNAi transcriptomes.

**Supplementary Figure S3.**
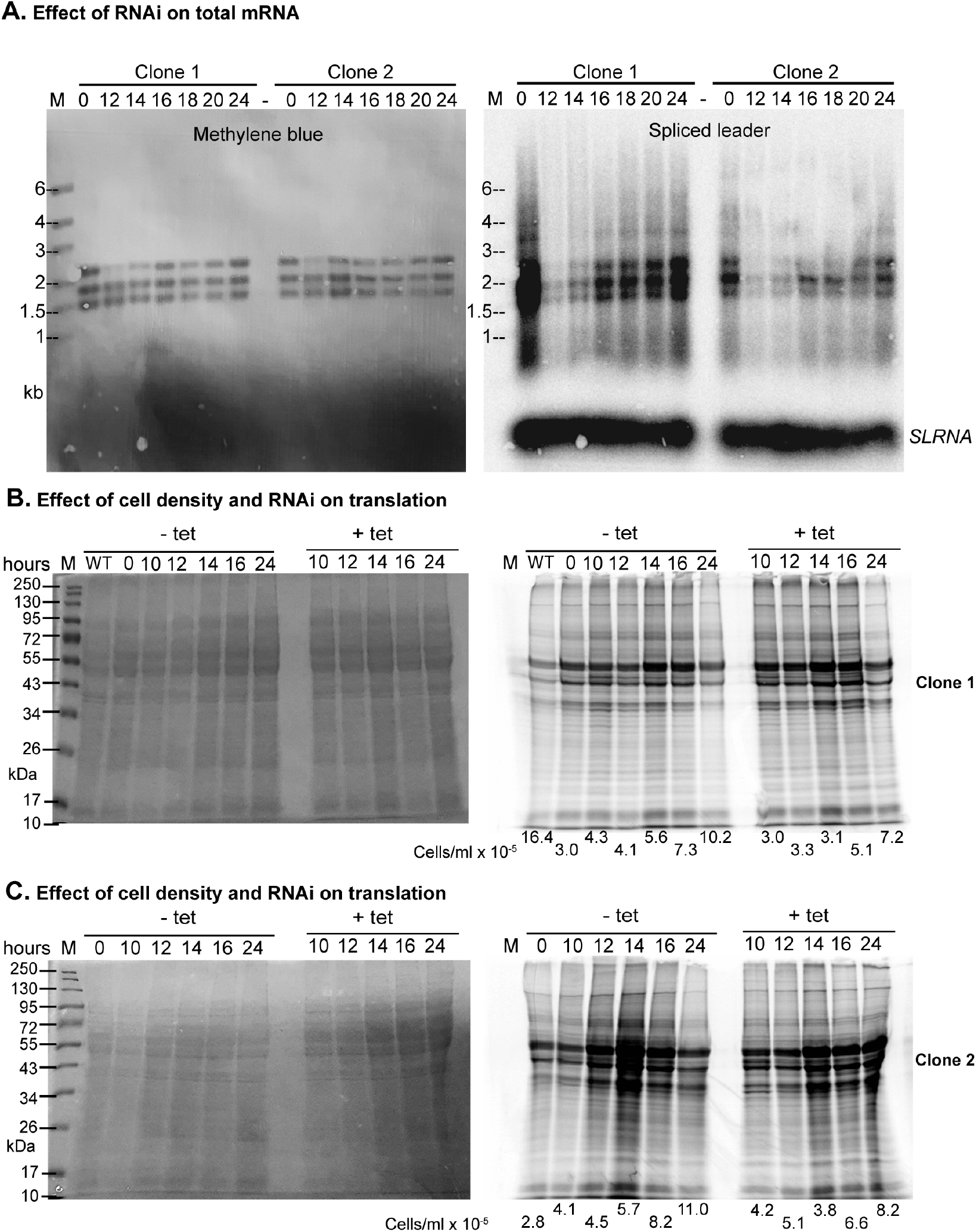
Effect of RNAi and cell density on mRNA levels and protein synthesis. A. Effect of ZC3H28 RNAi on total mRNA, detected with a spliced leader probe. There is some cross-hybridization with the rRNA but the signal above and below gives an indication of overall mRNA levels.. B. Effect of cell density and ZC3H28 RNAi on protein translation in Clone 1. The stained membrane is on the right and the 35S-Met+Cys incorporation on the right. Quantitation (using the protein stain as the control) revealed that the signal from radioactive incorporation was approximately halved at cell densities over 10^6^/ml. C. As for (B) but using clone 2. Quantitation was not possible because the protein stain was not even across the gel.

**Supplementary Table S1**

Complete mass spectrometry results. Details are on the first sheet.

**Supplementary Table S2**

RNAs associated with ZC3H28. Details are on the first sheet.

**Supplementary Table S3**

Effect of ZC3H28 on the transcriptome: results for all genes. Details are on the first sheet.

**Supplementary Table S4**

Effect of ZC3H28 on the transcriptome: results for the unique gene set. Details are on the first sheet.

**Supplementary Table S5**

List of plasmids and oligonucleotides

