## Supplementary Figure S1 for "Interactions of the *Trypanosoma brucei brucei* zinc-finger-domain protein ZC3H28"

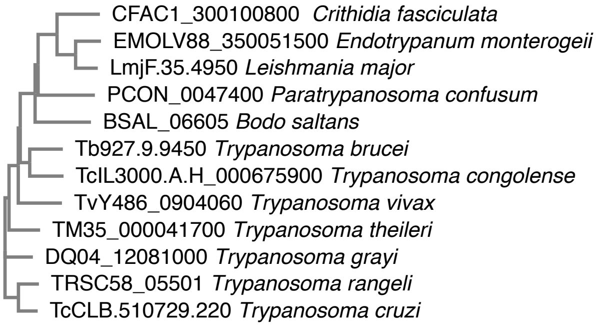


```
CFAC1_300100800             --MSASDSQS---------------------N--------------------GSPPLNRA     17
EMOLV88_350051500           --MSGLEAYGKAMG----------EVQSDRQM--------------------GSSSLTAA     28
LmjF.35.4950                --MSGFDSHGKAMG----------EAQSDRQM--------------------STSSLTAA     28
PCON_0047400                ------------------------------------------------------------     0
BSAL_06605                  --MSEPEQPLELAA--PPQPEVSTDAEVMNQEADATETPATASQPASIQTSASASPVADS     56
Tb927.9.9450                --MYSSEKEKEAPETLTSLPANAEEQQRER---DD-------N----HS-----------     33
TcIL3000.A.H_000675900      --MRS--SEKETPGAHILLPASVGEQQVEQ---DE-------G----QLQ----------     32
TvY486_0904060              MAMNLSGKEEWPDA-AVSPQ-----METEW---QA---------------GHDEAEVTA-     35
TM35_000041700              --MSSSDKERDTAERNMPPLNAADEKQVER---AA-------A----VAAGNGNDGNAN-     43
DQ04_12081000               --MSSSEKEREAQETGVSPSKPADENAVEP---AA-------------------------     30
TRSC58_05501                --MSSSEKEQDALKTGVPESSTAEDHEVRR---AA-------G----NGSGGEASTSAS-     43
TcCLB.510729.220            --MSSSEKEREAAENGVPTSDAAEEREVKQ---TA-------S----NGTGAEGSTATAA     44


CFAC1_300100800             SEEPRLTSSYLDGGANKTQPGNGGMGSDGTEHVPEVILHVRKNFGELAPVIEPYIYRSGY     77
EMOLV88_350051500           SR-TTPNPAPVDGGI-VSHVGGSPVDNDADEHVPPVIAHVRKNFGVLAPTIEPFIYRSGC     86
LmjF.35.4950                AR-ATPNPAAADGGA-VAHAGGSPVDNEMDEHVPPVIAHVRKNFGVLAPTIEPFIYRSGC     86
PCON_0047400                ------------------------------------------------------MFRLGW     6
BSAL_06605                  LPQPTPPPTP------SSAPPVPEASPVAAHHTPEVIAHLQEKFGLLSKTLEREVFRKGM     110
Tb927.9.9450                ------------------ASAQ-GEEDKNQDNTPVALLHLRQNFGVLTRVLEREVFRRGL     74
TcIL3000.A.H_000675900      ----------------AGAPARGEEEGKSNENVPPVLAHLQQNFGVMSRVLEREVYRRGL     76
TvY486_0904060              ---------S------SQVPHVVENGEEQHDNEPETVAHLRRNFGVLSRVLEREVFRRGL     80
TM35_000041700              -GNADGNDNR------GDSGDGPVAEEENNDHTPAVLLHIRQNFGVMSRVLEREVFRRGL     96
DQ04_12081000               -------------------SENPAAEEENQDRRLVTLVHLQQNFGVMSRVLEREVFRRGL     71
TRSC58_05501                ---LPQRQQQ------QSSGDGASAEEDSQDHRPVTLAHIQQNFGVMARVLEREVFRRGL     94
TcCLB.510729.220            TAAPLSQQKQ------QLSGDSAPAEEESQDHRPATLAHIQQNFGVMSRVLEREVFRRGL     98
                                                                                  ::* *


CFAC1_300100800             INGSPADVYQHLSRNYTFIVESALKQVGYEKFMMQMQVMLHHILYPEFLGDTDLLLNAIL     137
EMOLV88_350051500           LNKSVVDVYNHLSRSYPAIVDAALKQVGFEEFTMQMQIMLTQILSEEFGDDTEYLVNTIL     146
LmjF.35.4950                LNKSVVDVYNNLSRNYIAIVDAALKQVGFEEFTMQMQIMLTQILSEEFGEDTEYLVNTIL     146
PCON_0047400                INLTVSDLYGVLQANLNFLVDTMVRENGFQTVAWVLGEMMVLTLQPTCGEDAQSIVQAIQ     66
BSAL_06605                  INATVDDMYLRFEGQYQDLLNKVVSELSADVVLEELGNILHRSLLSTYNDSTPAIVVALQ     170
Tb927.9.9450                INSTVEDMYSAFSGASVVLVDILRSEVEPRLLRSQLGDMLYSNLELSYGAEHARMLATVI     134
TcIL3000.A.H_000675900      VQSTVEEMCQVFSTASMVLVDILRSEIEHRLLRSQMISMIYTALEASYGPDHARMLAMVI     136
TvY486_0904060              ISCTIPEMIEQFFIMSSPMYGIVKADMDQRVFHNQLFELLLTNLEQSYGAEHARFLASMI     140
TM35_000041700              INSSVEEMYSAFFLIAGLLIDILRSETDPRALRGQLVDMLFSIMEPNYGADHTRMLGAII     156
DQ04_12081000               INSSVEETYSVFFQVSSLLMEMFRMEIDPRVLRAQLLDMLLAILEPTYGADHTRMLMTII     131
TRSC58_05501                INSSVEEMYASFFVISTVLIEILKSETDQRLLHVQLLEMLLTILEPTYGIEHARMLVAIM     154
TcCLB.510729.220            INSSVEEMYSAFFVISGVLIEILKSETDQRLLHMQLLEMLLTILEPTYGIDHARMLVAIM     158
                            :. :  :    :      :      :     .   :  ::   :      .   ::  :


CFAC1_300100800             SIN-NHEALFVSLLSRSVLANLVNVAQQMIQNEMF----Y---SPQDGPSSGGGNRGMES     189
EMOLV88_350051500           TYT-GADSLFVSLLSKSMLDQVVTFAQSMMGGNMY----GMGYNMQGGGGGGSAGRGAEG     201
LmjF.35.4950                TYT-GADSLFVSLLSKSMLDQVITYAQSIMSGNMY----GMSYNMQSGGGGGSRGT---E     198
PCON_0047400                ENST-LYNQLYAVLNVDYLEEAVAEIQRSAVRR----VSGLRRMPPANNYAGS--RG-FS     118
BSAL_06605                  TAVLNLEGQFCAVINRQYLDAAVAMAVRTIERPQGQHRSHLRNS-QTDSPPDGMS-----     224
Tb927.9.9450                GSSLDDVYLFHAICSPSALEALVHQAQELINPPSNRTGGS-------NTYSDGA------     181
TcIL3000.A.H_000675900      DNSFDDLYLFNAVLSPTTLDSLVQQTQEWMNTSSSRAGSS-------STYNDSV------     183
TvY486_0904060              EQTVDEFYVFQALFSQHALVALVQQAEEYMNPQLRASSSS-------N-YGEGV------     186
TM35_000041700              EQVADDVFLFQAILSSAALDALVQRAQGYMNPQTSRSVGA-------G-ISGGVSGN-YS     207
DQ04_12081000               ERSANDLFLFQAILSSTSLDALVQRVQEIMNPP-ARSSGS-------N-MSGGVSGN-YN     181
TRSC58_05501                DQSVSDLYLFQALLSTSSLDALVQRAEEFMNPP-TRSVGA-------G-MSGGISGN-YS     204
TcCLB.510729.220            DQSVSDLYLFQALLSSSSLEALVQRAEEFMNPP-TRSVGA-------G-MSGGIPGN-YS     208
                                     : :: .   *   :                        .    .
```

```
CFAC1_300100800            GGGGGYYRGG-------------------------------------GGGGNNAGAQQGG   212
EMOLV88_350051500          GGGSNYYRGA-------------------------------------Q--------QQQG   216
LmjF.35.4950               GVGNSYYRGA-------------------------------------Q--------QHHG   213
PCON_0047400               SPPSSFAHGAPAPFGSPHHHHHHHLQQQQQQPVPVPVPAMVPSARGANPGREDAMDAVGG   178
BSAL_06605                 --HASDIRGD--------------IC-LDYEN-NRCT-----------------------   243
Tb927.9.9450               --------RA--------------VQSQSY------------------------GNAVGL   195
TcIL3000.A.H_000675900     --------RGG-------------AQAQSY------------------------GNTSGT   198
TvY486_0904060             --------RG--------------VQ-QQPSS-Y--------------------VDAAGP   202
TM35_000041700             GVPNENVRG---------------AQ-TSYNT-PANTNTMAT--------ANASSNTAAT   242
DQ04_12081000              SLPTENVRG---------------LQ-LGYNN-P-----------------------PAA   201
TRSC58_05501               SVSNENVRG---------------VQ-PTYNN-A-----------------------SAA   224
TcCLB.510729.220           GVQNENMRG---------------VQ-PAYNS-V-----------------------SAT   228
                                  :.

CFAC1_300100800            YRGG-ANANMRDH----------RGSMHSSINNHTHNH----ASNGNAGMAQSQ------   251
EMOLV88_350051500          YRNVGENGAPRDP----------RGG--------------------SVAVGLPH------   240
LmjF.35.4950               YRGVGENGSSRDL----------RGS--------------------SVAASPPH------   237
PCON_0047400               YRGPALS--MERAASTPSAMPVQPTPMMGGAAPAAVEGGGSAPRPSALAHSVPTGSATPT   236
BSAL_06605                 -RGANCR--YRHE----------GGPMEEAASP---------NRAGSHSMSIARGP----   277
Tb927.9.9450               ARPSFTE--FGKE----------RR-GE--VQP-------------TVEAKLPRVS----   223
TcIL3000.A.H_000675900     ARPSFSE--FGKD----------RR-VD--IQH---------Q--TGVDGMLPRGP----   228
TvY486_0904060             MRPSYTE--FVKE----------RR-GV--QQP---------HK-QINEPKLARGN----   233
TM35_000041700             VRPSYAD--YYKE----------RR-ME-----------------PSGDLKIVRGP----   268
DQ04_12081000              VRLPYGD--FSKE----------RR-GE-----------------QLGEPNVPRNA----   227
TRSC58_05501               VRPSYAE--FSKD----------RR-ME-----------------PALEPAMPRGA----   250
TcCLB.510729.220           ARPSYAE--FNKD----------RR-VE-----------------PSAEPAMSRGA----   254
                            *

CFAC1_300100800            -------ADGVGAQRPYRSTGMPGQAGNDAGGARERSSEDIRVGGDVLSQREESREWATS   304
EMOLV88_350051500          -------HDGVAPQKPYRSSGLPPQSS---DAVRERSSDDVRANSDALSQREEGREWSGG   290
LmjF.35.4950               -------QDGSTPQKPYRSSGLPPHSS---DPGRERSSDDLRATTDVISQREESRDWGGG   287
PCON_0047400               APSGAPGALAGANAAPYGTGGSLPLPPATTGPQGSH--------------------GLAG   276
BSAL_06605                 --------------LNQQ---APPA-----------------------------------   285
Tb927.9.9450               --------------NTYADGAAPAG-----------------------------------   234
TcIL3000.A.H_000675900     --------------AAYGDSVPAGG-----------------------------------   239
TvY486_0904060             --------------TPYVESSSPAQ-----------------------------------   244
TM35_000041700             --------------NTYTESNNAPV-----------------------------------   279
DQ04_12081000              --------------NAYTEGSNAPA-----------------------------------   238
TRSC58_05501               --------------NTYTEGGGAPA-----------------------------------   261
TcCLB.510729.220           --------------NTYTEGGGAPA-----------------------------------   265

CFAC1_300100800            QRAPPTPNPS--------------------------AQVDPTPSMRG------APPSSAT   332
EMOLV88_350051500          AVLPRSATVTGGG-----------------------VRVDSSLGPRGAASMISGMGAGAM   327
LmjF.35.4950               ASVPPRGATVTGG-----------------------VHIDSSVGPRGGAAPMSGMGAGAA   324
PCON_0047400               FQDPPVPHPSVGLHTSPPMHHQPPPPTSQAPQLPAPLQPQPQVVVGSASTLESGPTSGSD   336
BSAL_06605                 --------------------------ETSSV-----FGARPMVVLPQ-------------   301
Tb927.9.9450               --------------------------MNQPE-----EEVIPSVTTGG-------------   250
TcIL3000.A.H_000675900     --------------------------INQEG-----DAMLPA-VSGG-------------   254
TvY486_0904060             --------------------------ANSQD-----DDAVPA-NATG-------------   259
TM35_000041700             --------------------------SSQPE-----EDVMPM-ANSG-------------   294
DQ04_12081000              --------------------------SSQPE-----DDAMPT-VSSG-------------   253
TRSC58_05501               --------------------------SSQTD-----DEPTST-VTSG-------------   276
TcCLB.510729.220           --------------------------SSQPE-----DDAMPA-VASG-------------   280

CFAC1_300100800            SRQD------MAGSRP----QSEVPATAAVMPRHNHFIS--------------------P   362
EMOLV88_350051500          SRTD------MSGSRA----INEGAPGSGAMPRHTHHLF--------------------S   357
LmjF.35.4950               SRTE------VSVSRA----MSETASGAGAVPRHTHHHF--------------------S   354
PCON_0047400               SVSTFKPAPHITASRRQPGSSSAPPVAAGILLHHHHHHHLSLHL------HHHHIHHHQQ   390
BSAL_06605                 -----------AQSHHHHQIQLPSQAPPQSSVHHHHHLQSPLQVQHHLHHHHHHAQSTLQ   350
Tb927.9.9450               ----------WSNAQRKH-VEAE----REVLPHHA---------------HHHQLRARLP   280
TcIL3000.A.H_000675900     ----------WTNPQRRH-VEPE----REVPLHHG---------------HNHQLRSRPP   284
TvY486_0904060             ----------WSNSQRKH-VDHE----REVPVPQHHYHHHHHHH---HHHHNLQVHNRGS   301
TM35_000041700             ----------WVNAQRKQLADHD----RDASVHLQQQ-----------HHHHHHIHHRVP   329
DQ04_12081000              ----------WGNAQRKY-TDHD----RDVPIHHH---------------HHHLHHHRLP   283
TRSC58_05501               ----------WANVQRKH-AEHD----HDVPVHHYHQQQ---------QQQPHHIQHRVP   312
TcCLB.510729.220           ----------WAAVQRKH-TDLD----RDVTVHHHHHHQQQQQQ---QPQQQQHIQHRVP   322
                                         :
```

```
CFAC1_300100800             QNTAAA-TV-SVGKAAAAVERNVMS--GMPQHHFLPQRAATGAAMSRLPTPA--------    410
EMOLV88_350051500           SSTAVS----APSAKSSSMERAGIN--VVPQHNFLNSRTSPATNQARVTTPG--------    403
LmjF.35.4950                SAAAAAPAA-ALAAKLSSMERGGMT--GVPQHNLLPPRASPASSQPRVPTPG--------    403
PCON_0047400                QNADSGKPLHAIPTPL---PAQPL---PQPQSPQPQTQPSTLSLQPQQQSQPL-------    437
BSAL_06605                  ---A--PQS------------------QQHHHHHLQ------------------------    363
Tb927.9.9450                VNQFTAPQRPQV-------PSVPHHHHHHHHHHHVVVGTGPVGLQQQDR-----------    322
TcIL3000.A.H_000675900      MNQFTVPQRTHA-------PSVG----HHHHHHQVVVGTGSLGFQQQDR-----------    322
TvY486_0904060              LNQYAHQQRAQV-------PG-------VSH-HQVVVGSASVSLLQQPK-----------    335
TM35_000041700              FNQYP-QQRHQI-------PN-------VAHHHQVVVGSGPLGLQPQHQ-----------    363
DQ04_12081000               FNQYVHQQRHQL-------PN-------VAQNHQVVVGSGPMGLQPPQPQQHQQQQQQQQ    329
TRSC58_05501                FNQYLHQQRHQA-------PN-------VAHHHQVVVGAGPVALQQQQQ--------QQQ    350
TcCLB.510729.220            FNQYLHQQRNQA-------PN-------VAHHHQVVVGTGPMGLQQQQQ--------QQ-    359
                                                          :

CFAC1_300100800             ---STAASQNGS--PSPVPTVVRVGGNAMMHPHLHIGVPQQHQHQQQQQQMNNGGGA---    462
EMOLV88_350051500           ---VATVSLTSA--NSSMPTIGRGGPSIPMHHHH---------HHVGLGAMATSNAT---    446
LmjF.35.4950                ---APSVSQATT--TSPMPTMGRGGPSIPMHHHH---------HHLGIGAMATNSSA---    446
PCON_0047400                ---QQSQNQAPP----PRLSPSSGGNQHFPHHHHHLIPPQVHFAAHQQGQVPSVAGAAS-    489
BSAL_06605                  ------QPSGSPISASHNSPASFSPSQ-------------V-----VVGRVPAGSSMASV    399
Tb927.9.9450                HHQQLENPQNDAF---PRIPVATGLPK-------------R-----PVDRVLHQQSL---    358
TcIL3000.A.H_000675900      QQQQQDIVQTDTF---PRAPVPHGLSK-------------R-----PMDRLQHQQAM---    358
TvY486_0904060              QQQMQDAPQGESF---PRVPLSPAPSM-------------R-----MMDRISQ--GV---    369
TM35_000041700              -QQHQENPQGEPF---PRLPLPQGPHK-------------R-----VMDRMQL--NT---    396
DQ04_12081000               QQQQPELSQGGPY---PRAPVSQGPFK-------------R-----VGDRVPP--NA---    363
TRSC58_05501                QQQQQENHQTDSF---PRAPMPQGQQK-------------R-----MMDRLSP--GT---    384
TcCLB.510729.220            -QQQQESPQTDSF---PRAPVPQGQQK-------------R-----MVERLPP--GT---    392
                                                                              :

CFAC1_300100800             ---------------QGDWQGNNGNGGNMSPITPVTPQ----------RTVTPPSPQHMQ    497
EMOLV88_350051500           ---------------TSDWAAGNGASGSAPNMG-SIAQ----------RTLTPPTPQSAP    480
LmjF.35.4950                ---------------ASDWAASNSVSGSTTNIN-ASGQ----------RTLTPPTPQSAS    480
PCON_0047400                ---DNGTRAPPPARAPSLFQQQQPQPSQP--SLQAQPTPQVLSKSAPVAAPTPLQPHHQH    544
BSAL_06605                  KVLTQGATGTP----PSTFPSAQRSSPPP-SGFGGIPQPSQYHSG---SHPMTPPPPGSG    451
Tb927.9.9450                ----QGHDEASHQR-G-AWRGMGPSPTPQ-QGAAGVSQPNYNVRN---N-SPPVPSHVNH    407
TcIL3000.A.H_000675900      ----QGGEDAMHAR-G-VWRGASSSPAPQQQGAPGIPQASYNVRN---N-TSPLPPHATH    408
TvY486_0904060              ----QGPDD-VHVR-A-PWRGPLSSPGSQ-TGGAGVSQPSYNVTN---S-SPSPPVSHQG    417
TM35_000041700              ----QEPDE-PHSR-GLVWPGAPSSTGPQ-QDSMTVPQPSYHTPN---S-PPTPTPHLNH    445
DQ04_12081000               ----PEPDD-PHAH-GAMWRGAVPPSGPP-LEPVAVPQPSYHPQN---S-PPTPHSSHNP    412
TRSC58_05501                ----QESED-SHAR-GTMWRGSLPPPGAQ-HDGVTVSQSSYHSQN---P-SSPPTLH--T    431
TcCLB.510729.220            ----QESDE-PHVR-GPLWRVPPAPSGTQ-QETVTVPQPSYHAQN---P-SSTPTPH--S    439
                                              :

CFAC1_300100800             Q--------------QQQQ-------------------Q-QQQQPTRTMNIPAPLSVQRR    523
EMOLV88_350051500           V--------------QQHQLQ--------------------QQHPARTMNIPAPMSVQRR    506
LmjF.35.4950                M--------------QQHQHQH---M----QQQSQQQLP-QLQQPTRTMNIPAPLSVQRR    518
PCON_0047400                QIHTQHSHVHHHT----------QVPQSGLSQAQQPQLS-----LPP--KVVAPLPVTSG    587
BSAL_06605                  PLHHVHHHAHHSLGMHQHQHHHHHHLSAGLPQHQQQQYQQQQQQQQQ--QQVFP------    503
Tb927.9.9450                ------------------------------------------NQHPH--HIGPT------    417
TcIL3000.A.H_000675900      ------------------------------------------GQPLH--HMVTP------    418
TvY486_0904060              --H---------------------------------------GQQQH--GVGST------    428
TM35_000041700              --HPQHHHHHHHLLQHPHQIH--HQLQHQHQLQHQHQQQQQQHQQHH--QIPSP------    493
DQ04_12081000               --------------------------------------------HHH--QVVSP------    420
TRSC58_05501                --------------------------------------------NQH--QLPTP------    439
TcCLB.510729.220            --------------------------------------------NQH--QAASP------    447

CFAC1_300100800             A-----------------------------------------------------------    524
EMOLV88_350051500           T-----------------------------------------------------------    507
LmjF.35.4950                T-----------------------------------------------------------    519
PCON_0047400                STAPLMRSTPPPAPSVTAPTQIGHPPSVQQQSQGSYQPSNPPSTSPSPFPQGQPHSLQQQ    647
BSAL_06605                  --------------------------------------SAPQSSMGHVLPHQHHQHLHQH    525

CFAC1_300100800             ---------------------------------------------EERNA---------A    530
EMOLV88_350051500           ---------------------------------------------EKNAA---------A    513
LmjF.35.4950                ---------------------------------------------EKNAA---------A    525
PCON_0047400                ALSQPTVNVGR------PMLSPPPQQLHRQVQVAL--QPSVGPQPTQAP---VV------    690
BSAL_06605                  HQHQHQHHQHQQHQHQHHQHLQHHQQLHNQHHHQHQHMSSLQQQPQHNNNSGFPQQHGGS    585
```

```
CFAC1_300100800         AAAAAAAAAAAAAAAASAA--------------------------------------SAV    552
EMOLV88_350051500       AAAAAAAA--TNAASVSA----------------------------------------NG    531
LmjF.35.4950            AAAAAAAAAAAAPAAAAA----------------------------------------NG    545
PCON_0047400            ------AAAGANPVLRSILPMHRHNLQIFPLDSTPVVPLQTAPVGESVGVAQGHAVTHRQ    744
BSAL_06605              HAFHNDNSTHTAPSHAPPMHHHHHVLHHHHSS-------HLSNQ-------------GNN    625
Tb927.9.9450            ----------GSPTSARVIAPQRHNANQYIQ---------HPTM-------------TKQ    445
TcIL3000.A.H_000675900  ----------NSPTGTRILGPQRHNANQYMQ---------LGAV-------------GKQ    446
TvY486_0904060          ----------NSPSSVRPFTQHRHNISQHVP---------PGSM-------------GKP    456
TM35_000041700          ----------NSPTRSRPLVQHRHSVTHFVP---------PGSM-------------SKP    521
DQ04_12081000           ----------NSPSHPRMLVQHRHTINHFFP---------PGSM-------------GKQ    448
TRSC58_05501            ----------NSPTHSRFMLHQRHVTTHLVP---------PGPM-------------GKQ    467
TcCLB.510729.220        ----------NSPTHPRFLIHHRHATTHFVP---------PGPV-------------GKQ    475

CFAC1_300100800         KA--PSNNST------------SANNVSAASSSITPIVSDNVPSGRST------------    586
EMOLV88_350051500       SA--PTNSLS-------------NNS-------STPVVDAGIAP----------------    553
LmjF.35.4950            SA--PANSSS-------------PNN-------SSAAVEGGVAP----------------    567
PCON_0047400            SLPQPPASHQPQLLSAAPQMTSPPPMPSQRQSPGTPLVSGGITIPQPKGATATGNAVGSA    804
BSAL_06605              HVLQPNPQVQQQQ-----------------------------------------------    638
Tb927.9.9450            PIPQPTPSMASQQP-----------------------------SPQIS------------    464
TcIL3000.A.H_000675900  PLAQAAAPMASQQA-----------------------------TTQVS------------    465
TvY486_0904060          TSLQPTAAPTGQHN------------------------------PTQT------------    474
TM35_000041700          LSPQSTAQVTPQLT------------------------------PQPP------------    539
DQ04_12081000           MAPQPTTQMTPQTM------------------------------PQPA------------    466
TRSC58_05501            ASPQSATQISPQLT------------------------------PHLT------------    485
TcCLB.510729.220        ASPQPATQISPQKP------------------------------PHSA------------    493

CFAC1_300100800         ---------------ASLPH-------TTASAPPTAATPATP-------AARPALTPPVH    617
EMOLV88_350051500       ---------------------------RTMSAP-LSMNTNVP-------APRG-AHSLAH    577
LmjF.35.4950            ---------------------------RTMSAP-LGGSANVP-------APRG-AQSLPH    591
PCON_0047400            STFVSGGINVSIASKAQFPPQPLARGPTAQPLPPHPMRDSQPVASPSPQTPTVTAAAVSS    864
BSAL_06605              -------------------------------------RLAGPL--------QPV---PQH    650
Tb927.9.9450            ----------------------------AMNTTAPATPAVGAVSPPPPSAQRPL---PHH    493
TcIL3000.A.H_000675900  ----------------------------NMIPNPTAVNTTGPLASPS-STQRTV---VHH    493
TvY486_0904060          ----------------------------PP--ANTNMIHSS-STPPPQNSQRIV---LPH    500
TM35_000041700          ----------------------------A--------------TTPLPNSQRVV---VHH    554
DQ04_12081000           ----------------------------T--------------AAPPPNSQRVV---VHH    481
TRSC58_05501            ----------------------------PQPVTTTASTAAT--AASISNPQRVM---VP-    511
TcCLB.510729.220        ----------------------------TMTTTTTATTAAT-ASASITNSQRVM---IPH    521

CFAC1_300100800         HVHHMPIVHHHH------------------------HHHARLTTPAPQTAPATHSPAPPQ    653
EMOLV88_350051500       HQHPMPVLHHHH------------------------HHRNFTAPPVAVTAA-VQPRSPTH    612
LmjF.35.4950            HQHHMPVSHHHH------------------------HHYHFVASPPAVTAA-VQPPSATH    626
PCON_0047400            VRPHIHHHHN--I------------------------APHPLASVPVLMPS--QPSEGMY    896
BSAL_06605              HQHHLQHHHHQHQH---QHHAQHQHHHHHQHQHQHQHHHHQLHQPHHTVPHHLS-----H    702
Tb927.9.9450            HHH-LHLIQHHQQSQQNQQQQQQPHHHHHHHHHIH-HHHHDVMQQQPIVPPSPPPPQPSY    551
TcIL3000.A.H_000675900  ----LHMHHH-----QNQHQHHQQHPLPHHHHHIH-HHHHDLLQQQPVAPPAAVQAKHVY    543
TvY486_0904060          HHHHLHHHHHHHHH-----HHHH--HHHHQQQQQQ-QQQHDLPMQQPMTPSVVSPPQSMH    552
TM35_000041700          HHHHHLLHQQHHHH-----HHQQQQQQHQQQQQQS-PLPHELPLQHSIMPRVSTPPHSSY    608
DQ04_12081000           HH---HHHHHHFHH------------------------HHDLSLQQPGIPRVSPPPHTNL    514
TRSC58_05501            RH---HHH---HHH------------------------HHDIQPSQLMMSRASPPSQSIQ    541
TcCLB.510729.220        HH---HHH---HLH------------------------HHDIPPPQLMMPRVSPPSQSVH    551

CFAC1_300100800         QQQQ--------QQQQQTVEERAGSA-GTSIAATLNAGASLSSASDR-EG--SSSRTPAS    701
EMOLV88_350051500       SP--------------SASDERVAAIAADTINSTSQE-ANSS---PS-SA--TTSTLATA    651
LmjF.35.4950            LP--------------PASEERVATISADSTDSPVPE-TNLP---AN-QT--SAST----    661
PCON_0047400            FPVQSAPLPTQPHQQQPVVTP-QPHHQQSNTSASQPPGHVQPG--QQYQPPV---PTAT-    949
BSAL_06605              -HHQ------QHHQHHMQSAP-VLHTPNTAHSPSGSPHIPLPRSTAASHAPISIDAHDE-    753
Tb927.9.9450            -NVK------QEHTAAPETKP-MAH-NAE----AAPAQI-----KPLHHH--NVSFHPT-    590
TcIL3000.A.H_000675900  HNVK------QDHSATPETNA-IPH-NAE----VATSHV-----KPLHHH--NISIHST-    583
TvY486_0904060          LSVK------QEPATSPEATP-VAH-NVE----IAHPQL-----RPHFHH--QISFHHM-    592
TM35_000041700          QVMK------HEPIAPPEPKP-TTTHNVEAAAATTGAPI-----RPHFHH--HISAHPI-    653
DQ04_12081000           HAVK------QEAVAQPESKP-VAH-NVE----TAAPQL-----KPHFHH--QISFQHL-    554
TRSC58_05501            PLAK------PEAVNSPETKP-VAH-NAE----MAAPQL-----KPHHFH--HIPSAQK-    581
TcCLB.510729.220        AV-K------QEPVSPPETKP-VVH-NAE----TAVPQL-----RPHHFH--HIPPTQQ-    590
```

```
CFAC1_300100800          AAPAATVP-ATPTP-TPA-----------------------AVQPLRGMPRFPAHHH---   733
EMOLV88_350051500        ASTAASAA-ATKAPTMSA-----------------------PPSGLRGIGRIPIHHH---   684
LmjF.35.4950             SAPVRSPP-ATSASTTSA-----------------------PPPALRGLGRIPLHHH---   694
PCON_0047400             EMPAAAPS-PTPAPLAKP------------------AAVAPIPDVVRVAAPLPVHHHVLH   990
BSAL_06605               DMPLNRAP---STPQQP-----------------------------GSSPLA--AHHVHH   779
Tb927.9.9450             KLPQQPPP-QPSAPSSVG------------------------GEATHQVPIVPAHHHSIF   625
TcIL3000.A.H_000675900   KA----PP-QTNTPSSVG------------------------GEGAHHTPIVPTHHHSIF   614
TvY486_0904060           KMHQQAPL-RPNAPSSVG------------------------GDAVPHSTFVPTHHHSIH   627
TM35_000041700           RVPPPASP-QTNAPASV--GGGISVGVGGGGG--------ETSSHGGHGLMIPTHHHSIH   702
DQ04_12081000            KVLRPPTTQMN-APT---------------------SVGGVAAEPPSHGLMIPTHHHSIH   592
TRSC58_05501             VVRQPQPNVSVSASVGVGGGS-------GSGSGSGGGFSGGGSEAPSHGIMIPTHHHMLH   634
TcCLB.510729.220         KVLRQPQSVSVSASLGGGAGGGISTGSGGGGGGGGGGFNGGGSEAPSHGVMIPTHHHMIH   650
                                     :                                          **

CFAC1_300100800          --HHHGHLMSSRPDAAA-VAAAS---APPTPLPPAAAQQQQQQQAQQQQQQQHAQ--QQR   785
EMOLV88_350051500        --HHHIHLLVNRVETTM-PASSQR--QQQLPIPAASQPVSQPHLPPQPQRATTAIEVHEQ   739
LmjF.35.4950             --HHHMHLMVNKVDAPS-A---PQ--RQPQPIPATLQPQPQPHLPQPQQRVATAIDMHDQ   746
PCON_0047400             HHHHHHHHLLSHQMFHHHHHIQPMRPPHHHALPPHLAQGS----VSTQQQYTTAAPISID   1046
BSAL_06605               HVHHHHHALHHHHHVHHHHH------HHHH----QHQQAS----PQ--------------   811
Tb927.9.9450             HAQHVAPVHHHHHHHHHHH-------HHHH----HHLQQP----PHRYTSSPASAPMSVD   670
TcIL3000.A.H_000675900   QVQHTSSVHHHHHH----H-------HHHH----HLLQQQ----PYRYTSSSASAPMSVD   655
TvY486_0904060           QAH-IRAMG-----------------------------AQ----QHRHQAQTTTAPMSVD   653
TM35_000041700           HAQPIRAMMPLP----------------------QQQQQQ----QYRHPTSSAAAPMSID   736
DQ04_12081000            NFQQNRPVVPP---------------------------QH----QYRQSASSAPAPMSID   621
TRSC58_05501             HFQPIRQVVPSQ--------------------------QQ----QYRQLTSLTSGPMSID   664
TcCLB.510729.220         HFQQIRPSVPPP--------------------------QQ----QYRQPTTSAPGPMSID   680
                           :

CFAC1_300100800          AHAAAEEGPAQERAKEASGAW-PNEKEELRGALLGARVSEKGVTQFMTVDYGSVEELLMM   844
EMOLV88_350051500        HQFQQQRQQQQERKGTSNGRW-PNQKEELRGILLAARVSESAVNYLLTD--VNIDELRNL   796
LmjF.35.4950             --------QQQERTNAANGRW-PNQKEELRGMLLTARVSESAVTYLLTD--MNIEELRGL   795
PCON_0047400             AD---DEDMPLNR--ARTYSWRVAEREELDYFLEQHGLADEISSALR-SGTVSLNELRAM   1100
BSAL_06605               -----------------------RPAEDMSVFLSNAN-LTILVDPLS-KISITMEELRSL   846
Tb927.9.9450             MM---N-DEMTWA--LSSANLSPPPASELRRLLRERNVPESIVNSVL-EAGIKKEELLTM   723
TcIL3000.A.H_000675900   MI---NEDEMNWT--SKSAHSTVASAQELRCYLQDLGVQELLIDALV-KASVTKEELLSM   709
TvY486_0904060           AI---N-DDVSWS--TRLTNAPSSSVQELRCFLRDRSVPEDLINTVV-KAGFTKNELLEL   706
TM35_000041700           AL---SDEDMNWT--SSQATSMHSGVHELRCFLRNHMVPENVINALV-KTSLTLEDLMSI   790
DQ04_12081000            AL---SDEEMNWA--SPHAASMLPMTQELRCFFRDRSVPEAVINALV-KLSLTLDELLAI   675
TRSC58_05501             AL---SNDEMNWT--SSEAAAMLPGSQEIRCFLRERGVAEPVVNVLA-KMFLSLDELMAV   718
TcCLB.510729.220         AL---SNDEINWA--SSQ-SAMLSGPQELRYFLRERGVAETVINILA-KIFFSLDELMAA   733
                                                    ::   :            .      . ::*

CFAC1_300100800          KYVDFEKRLRPIVTGMLDRQRIRQLLMDNVARQSSPSENADSS-------------TTFS   891
EMOLV88_350051500        KYVDFEKKLRGGIPLLLERQRVRQALLDSATLQGSITDNDDSV-------------TSLS   843
LmjF.35.4950             KYVDFEKKLRGGIPLMLDRQRIRQVLSSSAAAQSQSSDNDDSA-------------TSSN   842
PCON_0047400             SRSTIEERLGSYGVQPQSRARLWNALHDGASDAAELA------------EKPHGSLRSLT   1148
BSAL_06605               SRGHLELRIKKYVCSKSMREQLWESLHPDGNYPTTPRKDEPAAAASPGHLDPISQLKHFA   906
Tb927.9.9450             SREMYDERLRTYLGS-QQSQTLWAALHSNEKSQHDRD------------GHEMNDLRSFI   770
TcIL3000.A.H_000675900   QKNVFEERFKKTLIP-PQRDFLWSVLHPGEKTLYDKD------------GHEMNDLRSFI   756
TvY486_0904060           SREVFDERLGRDVAGRQHLHFLWSVLHPEEKGPYDKD------------MPEMNDLRNFI   754
TM35_000041700           PREIFDEHIKKELVSANHREFVWAILHPNEKPLSEKD------------GPEMNELRNFI   838
DQ04_12081000            PREMFEEQLKKEVVSVHSRQYVWSVLHPNEKPLSEKD------------GPELHELRNFI   723
TRSC58_05501             PREVLDERLKKEVLSFNNRQYLWAALHPNEKLPSEKD------------VLEMSELRNFI   766
TcCLB.510729.220         SREAFEERIKKEIVSVNNRQYIWTILHPSEKLPSEKD------------VPEMNELRNFI   781
                              : ::            :   *

CFAC1_300100800          QRSASGVAGEER--SGNAQSMRPPSISVAPPNPGPPSPHSSPTSKSGVQVPQPRTRMQQQ   949
EMOLV88_350051500        RKSTAISVGTEEEPNGSGFPRRSLTVSSGMQSPL-SPQQSSPTSKTGVQVPQPRTKMQQQ   902
LmjF.35.4950             RKTATSGVGAEEEAGGSGLPRRPPAISTGVQSPV-TSHQSSPISKAGVQVPQPRTKMQQQ   901
PCON_0047400             LAG-DAKVGVELDRS-----------------VKSVPQRMAASAKQGMQLPQPRTVQGRQ   1190
BSAL_06605               ANN-AIASFGTRSKQTE---------ATSPQ-QDLKNSANTQAVKTMLNLPQPRSGVPTV   955
Tb927.9.9450             SNN-SIQNNIAQEAE-----------------DENGSSKRGRAAKPALQVPQPRTTQSRQ   812
TcIL3000.A.H_000675900   HNN-SIHNNMSQEVE-----------------DEGAAAKRGRPAKTALQVPQPRTSQSRQ   798
TvY486_0904060           NNN-SIHNNMGAEAE-----------------EDNGVSRRGRQGKSALHVPQPRTSQVKQ   796
TM35_000041700           HNN-AIHNNMSQDGE-----------------EENSASRRSRPSKTVLQVPQPRTSQSKQ   880
DQ04_12081000            HNN-SIHNNMAPDGG---------------E-EDSAMSRRGRQTKAMLQVPQPRTSQSKQ   766
TRSC58_05501             HTN-SIHNNMSQEGE-----------------DENATAKGGRTSKTMLQVPQPRTSQGKL   808
TcCLB.510729.220         HNN-AIHNNMSQDGE-----------------DDTTTLKPGRPNKNILQVPQPRTSQGKQ   823
                                                                     *  :::****:
```

```
CFAC1_300100800             QQQGHQ--N----------PNG---SPVASP-----------------------TDASYE    971
EMOLV88_350051500           QQQQSQQSV----------SGG---SAMASP-----------------------TETGLA    926
LmjF.35.4950                Q-QQTQQSI----------NGG---SAVASP-----------------------ADASFS    924
PCON_0047400                QQQQQIIPQ---------YSGGTREPNVAVITFSGSTPAARDQILNDIAQYANIVDSGLK    1241
BSAL_06605                  SSPASAKKP---PTSAEDAALYDGKEVRALLWISGTQRSNLKVIRETVSYYGSVLDCGYL    1012
Tb927.9.9450                QQQQQQ----------QQQQSNKGVEHRMILWLSGVSSSEVNQVLKEVGKYGKVVKHGVS    862
TcIL3000.A.H_000675900      QQQQQQQQQQQQQQQQQQQHPGEGVEYRAVVWLSGVSGSDPGQIVEKVSKHGKVLQYDVL    858
TvY486_0904060              -------------------QPAKPSEYKAVVWLSGVNSLEIPSVLEKVAKYGKVVQHGMS    837
TM35_000041700              QQQQQQQQQ---QQQQQQQQSGKGGECKAVIWLSGVTTTQMPSVLEEVSKYGKVLQHGTS    937
DQ04_12081000               QQ----------------QQPGKGGEYKAVIWLSGVNSGNPHLVFEELSRYGKVLQHGPS    810
TRSC58_05501                ------------------PQSGKVGEHNAVIWLSSVRSWQIQSVLEELSKYGKVLQHGTS    850
TcCLB.510729.220            QQ--------------QQQQSGKVGECKGVIWLSSVRSWQIQSAVEELSKYGKVLQHGAS    869
                                                                                   . .

CFAC1_300100800             -----------SSELESSRRES-SVRGGN--------------------PVTYANASS--    997
EMOLV88_350051500           -----------FSEKETAKRDS-ATKTES--------------------STTASIAPTSP    954
LmjF.35.4950                -----------SPEKESDKRES-GTKTGS--------------------PTTASGAPPSP    952
PCON_0047400                NNSYSKGTDVFFVKISNYSRDILGVRNVA--------GCDVENVQLVQPMGSEADALDSR    1293
BSAL_06605                  ----PRSKDAVYVKFSAYNKELQQTKHIPTGPGRHSESFAVVDLTLYNHGHIDDGSVPAI    1068
Tb927.9.9450                ----QDKSDMMYFKLKDCKGDLSGMQKIG--------NYIVEECHRVPPGEAGDGTPPA-    909
TcIL3000.A.H_000675900      ----SENSDMMYFKLKDCKADFSTVRKIG--------TFNVEDYYRVSPRDIGDGVPQK-    905
TvY486_0904060              ----V-DNNLFFCKLKDVKNDIHRLYKIG--------NAVVEEFYRVTPRPVDDGEPPL-    883
TM35_000041700              ----SQRPNLVYCKMGDHKNDLQMVRKIG--------SAVLEEYYRVPPGDIADGIPAT-    984
DQ04_12081000               ----SQKTDLIYCKLGDHKNELQTMRKIG--------GAVVEEFYRVSPRDIGDGAPPT-    857
TRSC58_05501                ----SHGHDLVYVKMGEHKNELQMVRKIG--------SAVVEDYYRVSPGDVDDGAPPTP    898
TcCLB.510729.220            ----SQRPDLIYFKLGDHKSELQMMRKIG--------SAVVEDYYRVSPGDVDDGAPPT-    916
                                         :      :

CFAC1_300100800             -------------------------------SG-----TFQPNSSTHNS-----------    1010
EMOLV88_350051500           APTP-----T----------------PASTPSP-----APTPNNPFSSN-----------    977
LmjF.35.4950                VPT---------------------------STP-----APSPNSSYSPS-----------    969
PCON_0047400                TDDSIVPLDG-------ATDQKRT--SGASRHT-----QSS-DNS---------------    1323
BSAL_06605                  PES--LISEARSSSDRHATRALRG--AGGSLTGSPHHNIPQPNSSFGRRSVGEDSNAEKS    1124
Tb927.9.9450                -RGP-RPEEAEED----GTNKLRAK-PNEAASPYPQQQTRP-------------------    943
TcIL3000.A.H_000675900      -PRA-RTEEAGDD----STNKARNK-PAGSGQPYQQQQQQQ-------------------    939
TvY486_0904060              -SIG-RDEEGEEN----EARKQRSKTASQ-PQASSQQNQHQ-------------------    917
TM35_000041700              -PTS-RPDEAEDS----ATNKARH-KVSGNTHPYP---QQQ-------------------    1015
DQ04_12081000               -PTG-RSEEGENN----GSSKARH-KAGGSANPYPSPPQQQ-------------------    891
TRSC58_05501                TPTG-RSEEADDN----GASKILGQKSGGGLQAYP---QHH-------------------    931
TcCLB.510729.220            -PTG-RPEGADDN----GPSKILSQKAGSGMHAYH---QHH-------------------    948

CFAC1_300100800             ----------------NRADEMEESREEDQRRGA---------GGGRRGGRRGGTRGGGF    1045
EMOLV88_350051500           ----------------TRNEEPDEVREEEHRRG-----------SGRRGGKRGGMGRGSF    1010
LmjF.35.4950                ----------------TRNEDPEDSREEEQRRG-----------SGRRGGKRGGLGRGGF    1002
PCON_0047400                --------LRRQSGGAELDETMEYDEAPGPRRGAKRPFG--MSQGGGRGNPRRPVNRGSH    1373
BSAL_06605                  NSFAGGRRGTTDSYD-HNDDDDEEMDDDRPRRGGKRDSNMGSRRGGASGSRRGGRGSGSQ    1183
Tb927.9.9450                --TSARQQGPAGSQV-DEH--EEDGNLEDSRT---------HRK--GMRNIR-GRGRGAN    986
TcIL3000.A.H_000675900      ----QQQQTHSGSYM-EEQ--EEDGDMEGSRT---------HRK--GMRQVRGGRGRGPH    981
TvY486_0904060              ----DLSQSNRESFV-DEQEQGEELEAEDGRA---------PRK--GTRHVR-ARGRGGH    960
TM35_000041700              ----HQGNPHRSSFG-EGD--EEDIDAEEARN---------HRK--SLRHTR-GRGRGGH    1056
DQ04_12081000               ----QQQSLHRGGYG-EGD--EEDIDGDEGRG---------HRK--GVRHSR-GRGRGGH    932
TRSC58_05501                ----HSP-ANRGNYG-VVD--EEDMDGDEIRN---------HRKGSGVRHTR-GRGRGGH    973
TcCLB.510729.220            ----QQPSANRGGYG-VVD--EEDMDGDETRN---------FRKGTGVKHAR-GRGRGGH    991
                                                  :       *                    *     *

CFAC1_300100800             -----------NSNRVCKFHGTPEGCQYGDKCYYIHGK------    1072
EMOLV88_350051500           -----------TSSRVCRFYGTAEGCQFGDKCHYVHNK------    1037
LmjF.35.4950                -----------SSSRVCRFFGTAEGCQYGDKCHYMHSK------    1029
PCON_0047400                RE---DGAEHTQSQSRCRYFEN-NTCKHGNDCKFLHLTN-----    1408
BSAL_06605                  DD---DSWDHYASNTVCKYYRS-GACKRGDTCKFAHPPQ-----    1218
Tb927.9.9450                HGYHTHRHSEGTGQVMCRYFNK-GACKYGEQCPFQHPSKHPGRS    1029
TcIL3000.A.H_000675900      HSYHAHRHNDNNGQTVCRFFSK-GVCKFGSQCQFSHLTKNMGRS    1024
TvY486_0904060              HVYA-RGHPEGFPGGLCRFFNK-GHCKHGGNCQFVHPKGPSRSQ    1002
TM35_000041700              HSYHSHRHSDGSSQALCRFFTK-GTCRNGDQCPYIHPKANSSRS    1099
DQ04_12081000               HNYHPHRHSEGNNQVICKFFSK-GICKFGDQCQYFHPAKSSSRS    975
TRSC58_05501                HNYHPHRHSEGSHQTVCKFFSK-GTCKYGDRCQFFHPTKHSSRS    1016
TcCLB.510729.220            HNYHHHRHPEGGTQMVCRFFSK-GTCKYGEHCQYFHPPKNASRS    1034
                                            *::. .   *: *  * : *
```
